# Copper-sensitive OsSPL9 TF regulates expression of *indica* rice domestication-associated miRNAs and phenotypes

**DOI:** 10.1101/2025.03.27.645634

**Authors:** Steffi Raju, Chenna Swetha, Chitthavalli Y. Harshith, P.V. Shivaprasad

**Affiliations:** National Centre for Biological Sciences, Research, GKVK Campus, Bangalore 560 065, India; SASTRA University, Thirumalaisamudram, Thanjavur 613 401, India; University of Copenhagen, Frederiksberg Campus, Bulowsvej 21, Frederiksberg 1870, Denmark

**Keywords:** Rice domestication, Cu-sensing miRNAs, SPL9 transcription factor (TF), miRNA targets

## Abstract

Domestication of *indica* rice facilitated better harvest and yield, however the molecular mechanisms that drove multitude of the associated phenotypes is poorly understood. A few genetic and epigenetic mechanisms have been attributed to *indica* rice domestication; however, upstream regulators of these variations are unknown. Here, we identified a copper (Cu)-dependent regulatory module, involved in the regulation of OsSPL9 TF and two classes of RNAs under its control. Differential accumulation of Cu-associated micro(mi)RNAs and Cu-associated protein-coding RNAs were a major portion of the differences between wild (*Oryza nivara*) and cultivated rice lines. We identified OsSPL9 as an upstream regulator of these changes through genetic and molecular analysis as well as by using Cu stressed conditions. OsSPL9 bound to the promoters of these genes through a conserved GTAC enriched motif. OsSPL9, Cu-associated miRNAs and their targets acted as a regulatory loop, since mis-expression of SPL9 alone, or any individual Cu-associated miRNA, also altered levels of other Cu-miRNAs and their cognate targets. OsSPL9-mediated regulation was closely linked to Cu accumulation and metabolism, indicating previously unappreciated roles of metal ions in mediating domestication-associated phenotypes. Our study facilitates a better understanding of the crosstalk between genetic and epigenetic regulation that contributed to *indica* rice domestication.

## Introduction

Domestication is a phenomenon of evolution of crops assisted by cultivation and it has induced diverse changes in plants, often referred to as domestication-associated traits (Doebley 2006). Although genomic loci attributed to these phenotypes have been characterised to a greater extent from several studies in diverse crops, mechanisms underlying this process have not been well understood. This difficulty was largely attributed to the fact that not all phenotypic changes in the course of domestication could be attributed to protein-coding genes (Meyer and Purugganan 2013). Specifically, a majority of the QTLs attributed to distinct crop phenotypes did not overlap with specific protein-coding genes and this led to the hypothesis that small (s)RNAs or other epigenetic regulators might be playing a dominant role in establishing domestication-associated phenotypes (Shivaprasad et al. 2012a, 2012b; Houston et al. 2013; Song et al. 2017; Shivaprasad 2019). sRNAs as major regulators of gene expression and phenotypes in plants was well elucidated, along with its role in inducing heritable epigenetic marks such as DNA methylation and histone modifications (Hollick and Chandler 2001; Baulcombe 2004; Seymour et al. 2008; Niederhuth and Schmitz 2014). Indeed, several landmark studies have revealed that there is a major influence of sRNAs including miRNAs in regulating gene expression changes resulting in crop domestication phenotypes. In agreement with this, significant number of yield-related QTLs across crops overlapped with sRNAs, their targets or players involved in epigenetic modifications (Wang et al. 2010; Campo et al. 2013; Qin et al. 2014; Swetha et al. 2018).

The cultivated rice is thought to have originated from two wild relatives of cultivated rice, namely, *Oryza nivara* and *Oryza rufipogon* (Aggarwal et al. 1997). Among these, *indica* rice is of generally shorter stature with morphological and physiological features distinct from those of *japonica* and *javanica* rice lines. Several studies identified genetic and epigenetic regulators of *indica* rice domestication. Well-known phenotypes and associated genes identified in *indica* rice include increase in grain shape (Lu et al. 2013), grain colour, heading date (Takai et al. 2021), loss of shattering (Lin et al. 2007), cold tolerance (Wu et al. 2024), and thermo-tolerance (Wang et al. 2016). Several sequence variations including SNPs in coding as well as regulatory regions, transposon insertions and gene duplications appear to have contributed to *indica* rice domestication (Konishi et al. 2006; Huo et al. 2017; Shivaprasad 2019). Among the epigenetic factors, sRNAs and methylation have been documented to influence rice domestication phenotypes.

A major domestication associated phenotype determinant in *indica* rice is through miRNA-mediated laccase silencing (Swetha et al. 2018). The miR397 is a Cu-dependent miRNA (Panda and Sunkar 2015; Huang et al. 2021) and it can regulate the expression of Cu-binding proteins such as laccases post transcriptionally (Wang et al. 2014; Gaddam et al. 2024). More than 25 QTLs implicated in rice domestication overlapped with miR397 or laccases indicating its importance in the domestication process (Swetha et al. 2018). Several reports further indicated that miR397 levels can also reinforce phenotypes that are associated with yield, such as grain weight (Zhong et al. 2020), seed number (Wang et al. 2014), grain yield (Yu et al. 2024), seed size (Guo et al. 2024), and circadian rhythm (Feng et al. 2020). The role of miR397 to regulate these diverse phenotypes associated with yield was also conserved across other food crops such as wheat (Wang et al. 2024a). This indicates the potential role of miRNAs as emerging QTLs across food crops in regulating phenotypes associated with domestication.

The class of Cu-dependent miRNAs includes miRNAs such as miR397, miR528, miR408, and miR398 (Burkhead et al. 2009; Pilon 2011, 2017; Zhu et al. 2020). However, the role of miRNAs other than miR397 in the context of domestication is not understood. miR528 is a conserved monocot-specific miRNA (Chen et al. 2019; Zhu et al. 2020), which is known to regulate flowering time in rice (Yang et al. 2019), tillering phenotypes in switchgrass (Han et al. 2024), and broad-spectrum antiviral resistance in rice (Yao et al. 2019). miR408 has the potential to influence grain yield and photosynthesis in rice (Zhang et al. 2017), and might contribute to anthocyanin biosynthesis (Hu et al. 2023b). The role of miR398 was majorly studied in the context of abiotic stress responses in rice (Lu et al. 2010, 2022). How exactly Cu or these Cu-dependent miRNAs play a role in influencing these phenotypes is unknown.

Among the genes that regulate domestication-associated phenotypes are TFs or their cofactors (Doebley et al. 2006; Purugganan and Fuller 2009; Meyer and Purugganan 2013; Swinnen et al. 2016, 2019; Shivaprasad 2019). Interestingly, some or most of these Cu-dependent miRNAs are under the regulation of a Squamosa promoter-like TF, named SPL7 in *Arabidopsis* (Yamasaki et al. 2009; Araki et al. 2018) and at least one miRNA is under the control of OsSPL9 in rice (Tang et al. 2016; Yao et al. 2019, 2022; Hu et al. 2021; Wang et al. 2024b). Among Brassicales, SPL7 ensures Cu acquisition and localised deposition in seeds to enable its dispersal (Pérez-Antón et al. 2022), and also mediates the crosstalk between sucrose signalling, growth and copper homeostasis as documented in Arabidopsis (Ren and Tang 2012; Schulten et al. 2022). Rice OsSPL9 is known to regulate antiviral response through miR528-mediated post transcriptional gene silencing of ascorbate oxidase, rendering immunity (Yao et al. 2019), and also as a direct/indirect determinant of grain size and yield through unknown mechanisms (Hu et al., 2021).

The process of domestication in food crops led to major shifts in the nutritional requirements due to altered growth habitats and conditions (Li et al. 2024). It is well-established that dependency on metalloproteins acting as enzymes and other catalysts likely ensured better survival and fitness in life forms including crops (Pilon et al. 2009; Walker and Waters 2011; Yruela 2013). Allowing a permissible range of nutrient accumulation seems to be of utmost importance to balance out excessive metals to avoid mis-incorporation and toxicity within cells. Nutrient balancing during domestication was also essential to mediate phenotypic plasticity in plants over the course of selection eventually leading to altered phenotypes (Perkins and Lynch 2021). How metal ions such as Cu and other micronutrients and their upstream regulators might have emerged in a context where such nutrient dependency has evolved is not understood.

Here we show that, Cu-responsive miRNAs and their targets are a substantial portion of changes between cultivated and wild rice lines. Through genetic and molecular approaches, we show that OsSPL9 is the upstream regulator of these miRNAs and other differentially expressing genes between wild and cultivated lines. Using ChIP-seq, we identified that OsSPL9 binds to GTAC enriched motifs of Cu-associated genes. Several rice domestication-associated phenotypes under the control of SPL9 and miRNAs were altered under limiting and excess Cu conditions, indicating how Cu might have influenced these phenotypic changes. These results indicate potential contributions of altered metal ion availability and their accumulation influencing crop phenotypes in agricultural systems.

## Results

### Substantial changes in micronutrient related genes between wild and cultivated rice

In order to identify differentially expressing genes between wild and cultivated rice line Pusa Basmati 1 (henceforth PB-1) across various growth stages, we performed RNA-seq analysis and obtained a mapping efficiency of 95 % to rice genome (IRGSP-1.0) (Supplementary Table 1). Nearly 25 % of the genes that showed higher expression in *O. nivara* when compared to PB-1 were associated with disease resistance and secondary metabolic pathways (Supplementary Figure 1A, B). Such a relationship has been well-explained across several wild species and domesticated crops (Wang et al. 2015; Dong et al. 2017; Thein et al. 2019; Sinha et al. 2023).

Among the prominent genes that exhibited reduced expression in cultivated rice PB-1 included disease resistance-associated MYBs, WRKY, RALF, and bZIP TFs (Fig 1A lower panel). About 713 genes out of 1335 DEGs had higher expression in PB-1 (Fig 1A upper panel). As expected, growth-related and yield-associated genes such as *OsGLW7* (Si et al. 2016)*, OsCLV2c* (Shen et al. 2024) were upregulated in PB-1, along with some of the most prominent macro and micro-nutrient and metal ion-associated genes. It is well known that improved nitrate metabolism is a major change responsible for increased yield across cultivated lines (Gao et al. 2022) and in agreement with this, around 70 nitrate-responsive genes showed higher expression in PB-1 (Supplementary Figure 1C). Similarly, iron-responsive genes also exhibited higher expression in PB-1 importance of which is unknown (Supplementary Figure 1D). Additionally, the expression of several phosphate-responsive genes was high in wild rice when compared to PB-1 (Supplementary Figure 1E). For example, a central regulator of phosphate signalling in rice named OsPHR2 and a high affinity phosphate transporter-OsPT4 were upregulated 2-fold and 16-fold, respectively, in *O. nivara* (Figure 1B). As implicated previously, phosphate levels and expression of phosphate-responsive genes are important factors in disease resistance (Val-Torregrosa et al. 2022; Paries and Gutjahr 2023). Several genes associated with biosynthetic processes also showed a significantly higher expression in wild rice compared to the cultivated rice line (Supplementary Figure 1F).

**Figure 1.**
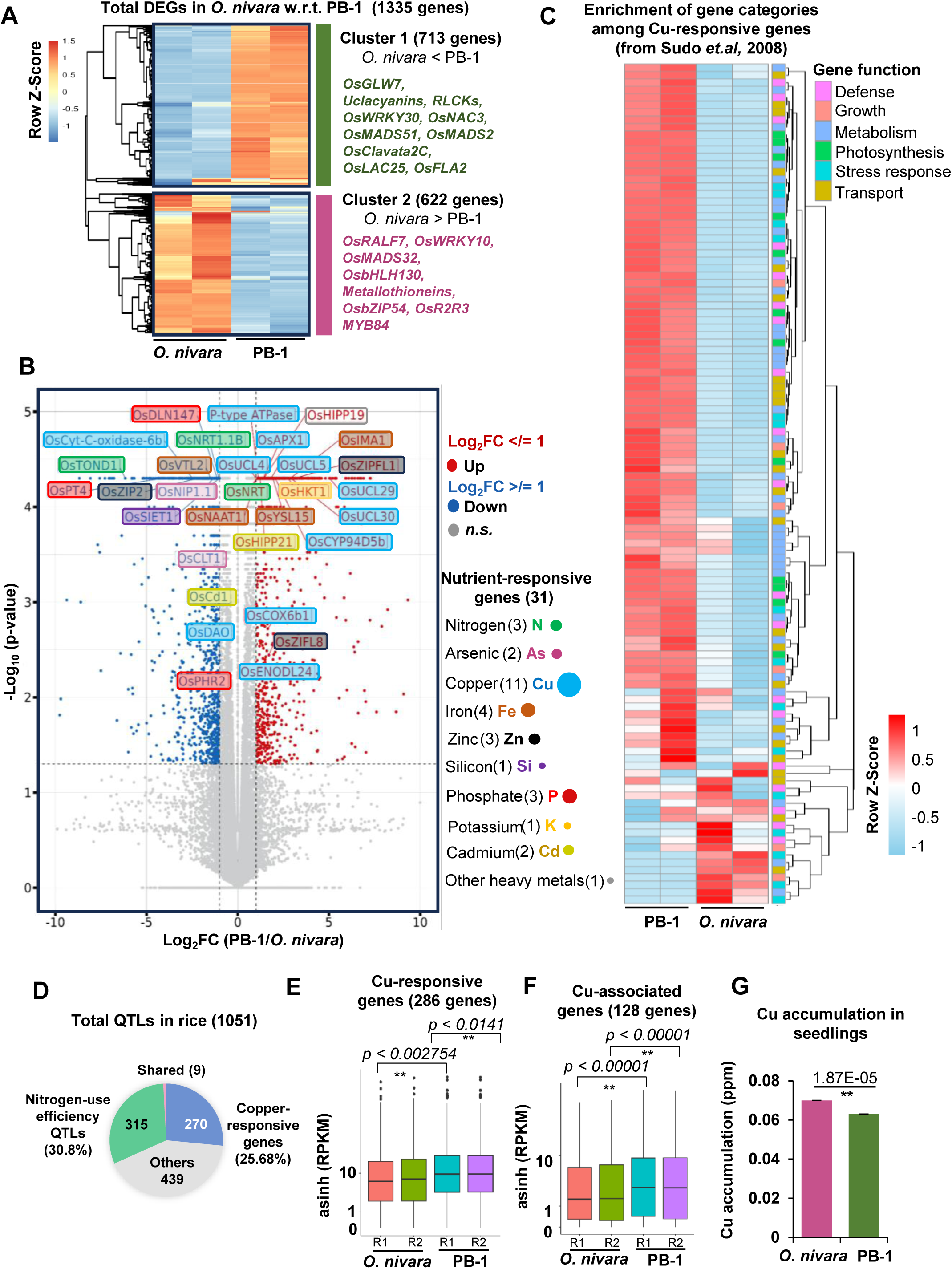
Wild and domesticated varieties show nutrient-associated DEGs and differential copper accumulation. (A) Clustered heatmap showing the differentially expressed genes between *O.nivara* and PB-1. Cluster I (713 genes) and II (622 genes) are marked with the representative genes of each cluster. The row Z-score of the FPKM is plotted for the heatmap. (B) Volcano plot showing the nutrient-responsive DEGs (seedlings) and their distributions. Blue dots-downregulated genes, red dots-upregulated genes, *n.s.*-non-significant. The coloured dots on the right indicate the respective nutrients and the associated genes among the DEGs. The size of the dot indicates the number of genes belonging to each category. (C) Annotated heatmap showing the gene functions of the Cu-responsive genes across *O. nivara* and PB-1. The row Z-score of the FPKM is plotted for the heatmap. Annotations of gene functions are indicated. (D) Pie-chart showing the distribution of nitrogen-use efficiency QTLs (324) and copper-responsive genes (286) among the 1051 QTLs in rice. (E) Box plot showing expression of Cu-responsive genes (286) in wild and cultivated rice seedlings. The y-axis is scaled to the inverse-sine hyerbolic function. (F) Box plot showing expression of Cu-associated genes (128) in wild and cultivated rice seedlings. The y-axis is scaled to the inverse-sine hyperbolic function. (E-F) asinh converted RPKM value were used for boxplot. The boxes show median values and interquartile range. Whiskers show minimum and maximum values. Comparisons were made with two-sided Wilcoxon test (*p*-value < 0.05 was considered significant). (G) Cu-accumulation in 21-day old seedlings of *O. nivara* and PB-1 was performed using ICP-OES. The concentration of Cu in parts per million (ppm) is plotted. An average of 3 technical replicates were used. Two-tailed Student’s t-test was used for statistical comparison. (*) *p*-value < 0.05, (*ns*) non-significant.

Among the genes involved in micronutrient and metal ion response, a major category of upregulated genes in cultivated lines were Cu-binding proteins such as uclacyanins (Figure 1B). Among the 6 categories of Cu-responsive genes classified based on functions in rice (Sudo et al. 2008), major categories of highly expressed genes in PB-1 included those involved in photosynthesis, metabolism and transport. Very few genes among Cu-responsive genes were upregulated in *O. nivara* and they included categories such as response to stress and defense as indicated earlier (Figure 1C). The expression levels of Cu-responsive genes (as listed in (Sudo et al. 2008) and about 128 Cu-associated genes (curated from RAPDB) were upregulated in cultivated rice when compared to *O. nivara* (Figure 1E-F). This indicates that Cu-responsive modules were among the major DEGs between *O. nivara* and PB-1.

Among these Cu-associated genes are a family of laccases that were previously implicated in *indica* rice domestication (Swetha et al. 2018). In agreement with the large number of Cu-associated DEGs identified between wild and cultivated rice, a large number of QTLs (from Gramene QTL database) for all traits also overlapped with Cu-responsive genes (25% of all QTLs). In comparison, among the 327 well-characterised QTLs for nitrogen-use efficiency (Kumari et al. 2021), there was an overlap of 30.8% (Figure 1D). These results indicate a previously unappreciated contribution of Cu in mediating domestication-associated changes.

We hypothesised that the observed differences in gene expression for Cu-responsive genes might be due to differential absorption/accumulation of Cu between wild and cultivated species. Several studies indicated altered accumulation of nutrients between wild and cultivated crops that resulted in above and below-ground growth phenotypes in wheat (Gioia et al. 2015). Recently, it was also reported in durum wheat that a 2-fold increase in DEGs occurred depending on nitrogen availability during domestication (Pieri et al. 2024). Surprisingly, levels of Cu measured through Inductively-Coupled Plasma-Optical Emission Spectroscopy (ICP-OES) showed higher accumulation in *O. nivara* seedlings when compared to cultivated line PB-1 (Figure 1G). This indicated a possible reason why the Cu-associated genes were among the DEGs. It is well-known that Cu in vegetative tissues is inhibitory to herbivores as well as other biotic stress factors (Liu et al. 2015a; Chai et al. 2020; Yao et al. 2022) and several studies suggest that certain hyperaccumulator plants ward off pathogen and pests by selective allocation of transition metals (De et al. 2025). Although several heavy metals such as arsenic are known to accumulate in rice grains (Yao et al. 2021), genes associated with arsenic were not enriched among the DEGs between the wild and cultivated lines. While these results are in agreement with general assumption that wild species are able to accumulate bivalent anions and cations, they do not provide insights why Cu-associated genes are among the most DEGs or which tissues copper accumulates between wild and cultivated lines.

### Cu-associated miRNAs and their targets are differentially expressed in wild and cultivated rice

miRNAs are well-known regulators of large family of genes and they fine-tune gene expression during development and stress responses (Baulcombe 2004, 2023; Shivaprasad et al. 2012a). Several domestication associated DEGs and phenotypes are regulated by miRNAs and other sRNAs in rice (Liu et al. 2015b, 2016; Yao et al. 2015; Tang and Chu 2017; Swetha et al. 2018; Shivaprasad 2019). Based on degradome analysis (Swetha et al. 2022), we identified several differentially expressing miRNAs between wild and cultivated rice that targeted genes such as Os5NG4 and R2R3MYB. In order to get a comprehensive view of nutrient-associated miRNAs and their target genes, we analysed sRNA datasets across different tissues of wild and cultivated rice lines. Several nutrient associated miRNAs (Budak et al. 2015) were unchanged in seedling, flag leaf and panicle tissues across wild and cultivated lines (Figure 2A). These include miR390 which is cadmium-responsive (Ding et al. 2016), miR395, sulfate-responsive miRNA (Panda and Sunkar 2015), miR156 responsive to phosphate and potassium (Lei et al. 2016), miR393 associated with nitrogen (Li et al. 2016), and miR168,171 and 172 that are associated with Fe accumulation (Kong and Yang 2010). The miRNAs that showed differential expression between wild and cultivated rice varieties were miR397, miR398, miR408 and miR528 across several different tissues such as seedling, flag leaf and panicles (Supplementary Figure 2A). These set of miRNAs are known to establish Cu homeostasis and are known to differentially express upon sensing Cu levels (Burkhead et al. 2009; Pilon 2017). These miRNAs majorly targeted genes encoding Cu-binding proteins (Pilon 2011, 2017).

**Figure 2.**
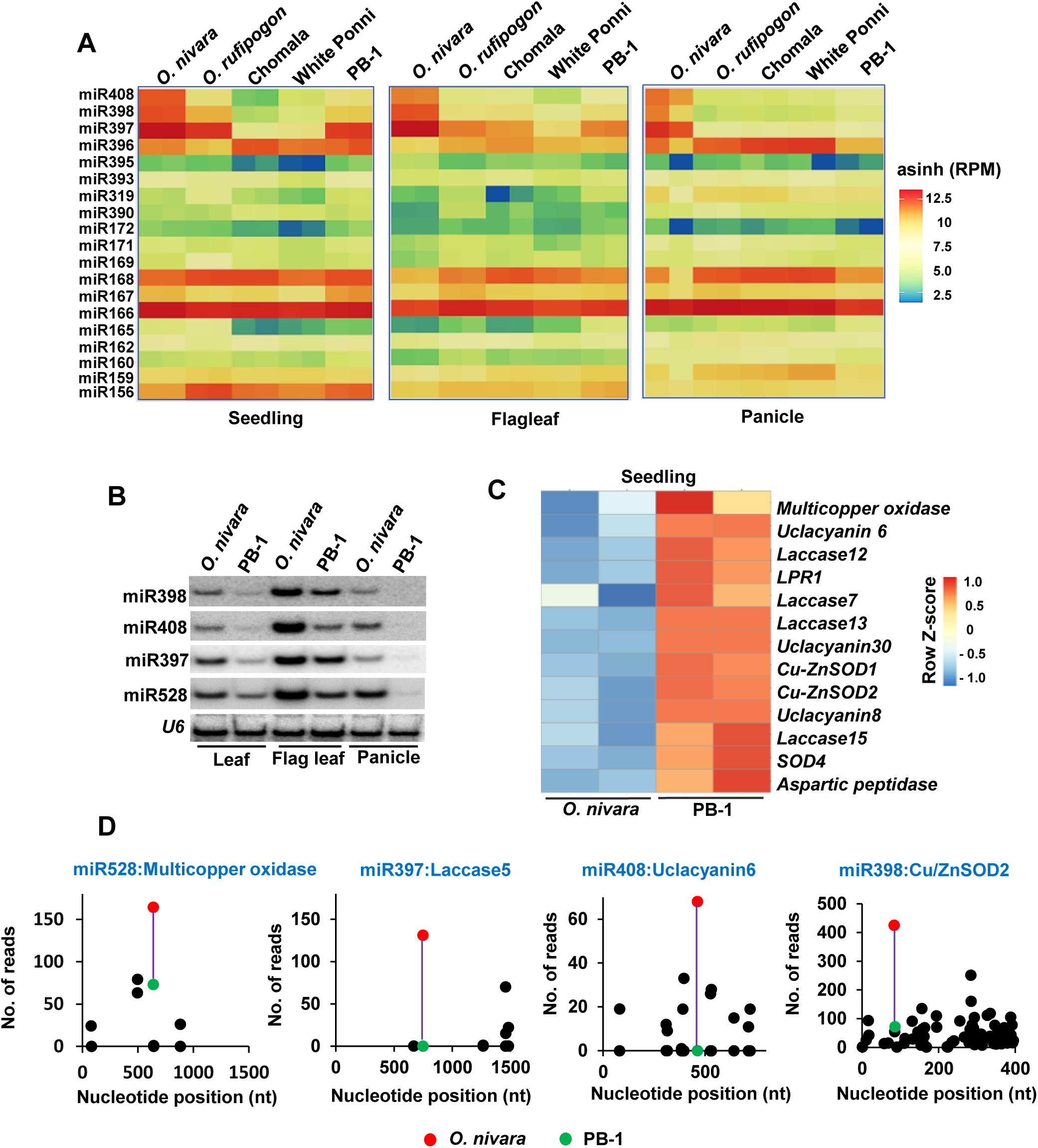
Differential accumulation and targeting of Cu-miRNAs in wild and cultivated rice. (A) Heatmap showing nutrient-responsive miRNAs among wild rice (*O. nivara* and *O. rufipogon*), cultivated lines (White ponni and PB-1) and a landrace (Chomala). The expression values are inverse-sine hyperbolic function of differences (Reads per million-RPM). (B) Northern analysis for miRNAs. *U6* serves as control for the miRNAs. (C) Heatmap showing the differences in the expression of miRNA targets in *O. nivara* and PB-1 seedlings. The row Z-score is plotted for the RPKM value of the samples. The genes represented are the targets of the Cu-responsive miRNAs. (D) Degradome analysis of the miRNAs and their targets. Red dots indicates reads from *O. nivara* and green dots indicate reads from PB-1. Purple line indicates the trend line in the degradome reads between the samples.

In addition to PB-1, another cultivated *indica* rice line White Ponni also showed patterns similar to PB-1 (Supplementary Figure 2A), indicating that the differences in miRNA levels are more general. Higher expression of Cu-miRNAs was also observed in *O. rufipogon*, ancestor of several *indica* and *japonica* accessions. For example, *O. sativa japonica* Nipponbare, a cultivated *japonica* rice, accumulates reduced Cu-associated miRNAs such as miR408 (5-fold reduction when compared to *O. rufipogon*), and miR528 (30-fold reduction when compared to *O. rufipogon*) (Supplementary Table 6; Data reanalysed from (Wang et al. 2012). However, another Cu-dependent miRNA, miR397 did not show a reduction between wild relatives - *O. rufipogon* and Nipponbare (Supplementary Table 6). Also, miR397b that is highly expressed in *O. nivara* is less abundant in *O. rufipogon*, indicating a significant difference between *indica* and *japonica* rice lines. This indicates that roles of Cu-dependent miRNAs in general and miR397-related domestication-associated phenotypes in particular, might not be entirely similar between *indica* and *japonica* rice lines. These differences between subspecies are interesting and it is possible that they might have evolved to match the locations/habitats in which they were domesticated.

To further confirm the differential accumulation of miRNAs, we performed a sRNA northern blotting that indicated high expression of Cu-miRNAs in all tissues of *O. nivara* (Figure 2B). We used psRNATarget tool (Dai and Zhao 2011; Dai et al. 2018) to identify the putative targets of these Cu-miRNAs and found additional, previously unknown, targets such as members of SOD and uclacyanin families that are also Cu-associated genes. Previously known as well as newly identified targets of these miRNAs also showed differential expression between wild and cultivated rice lines as seen in RNA-seq analysis (Figure 2C). The targets and miRNA levels showed negative correlation indicating the miRNA mediated regulation of these Cu-responsive target genes.

To further confirm differential targeting of these Cu-associated miRNAs, degradome sequencing (Swetha et al. 2022) was employed. As expected, targets of these miRNAs such as laccases, uclacyanins, multicopper-oxidases and Cu/Zn-SODs had higher degradome reads, often in category 0, in *O. nivara* when compared to PB-1, indicating a much stronger targeting in wild rice due to higher levels of miRNAs (Figure 2D). A miR528 target named multicopper-oxidase had 164 degradome reads from wild rice and 73 reads from cultivated rice. Similarly, Cu/Zn-SOD had 425 degradome reads in category 0 from *O. nivara* when compared to 121 reads of category 2 in PB-1 flag-leaf and panicle tissues. The targets of these miRNAs are known to regulate several yield-associated phenotypes, and these modules include miR397:laccases (Wang et al. 2014; Swetha et al. 2018; Zhong et al. 2020; Guo et al. 2024; Yu et al. 2024), miR528: RFI2 (Yang et al. 2019), miR408:Uclacyanins (Zhang et al. 2017), and miR398:Cu/Zn-SOD (Lu et al. 2010) (Supplementary Figure 2B).

### OsSPL9 is an SBP-domain containing TF upstream of domestication-associated miRNAs

miRNA-TF associated networks are well studied in plants (Wu and Poethig 2006; Poethig 2009; Yamaguchi et al. 2009; Xu et al. 2016). TFs associated with basal metabolism and development remain conserved across land plants but several nutrient related TFs were co-opted with the downstream gene regulatory networks to elicit specific responses in changing environmental conditions (Dong et al. 2024). In Arabidopsis, the Cu-responsive miRNAs (miR397, miR398, miR408, miR857) are regulated by the SBP-domain containing TF named *AtSPL7* (Yamasaki et al. 2009). To identify rice homologue of *AtSPL7*, we performed a phylogenetic analysis for SPL family of genes in Arabidopsis and rice (Figure 3A), having 16 and 19 members, respectively (Preston and Hileman 2013). This analysis indicated that OsSPL9 is homologous to *AtSPL7*. Interestingly, OsSPL9 in rice was identified as a regulator of miR528 (Yao et al. 2019). In agreement with a possible role for OsSPL9 as an upstream regulator of Cu-associated miRNAs, Cu-miRNA precursor levels were reduced in specific tissues of *Osspl9ko* lines in *japonica* rice (Wang et al. 2024b).

**Figure 3.**
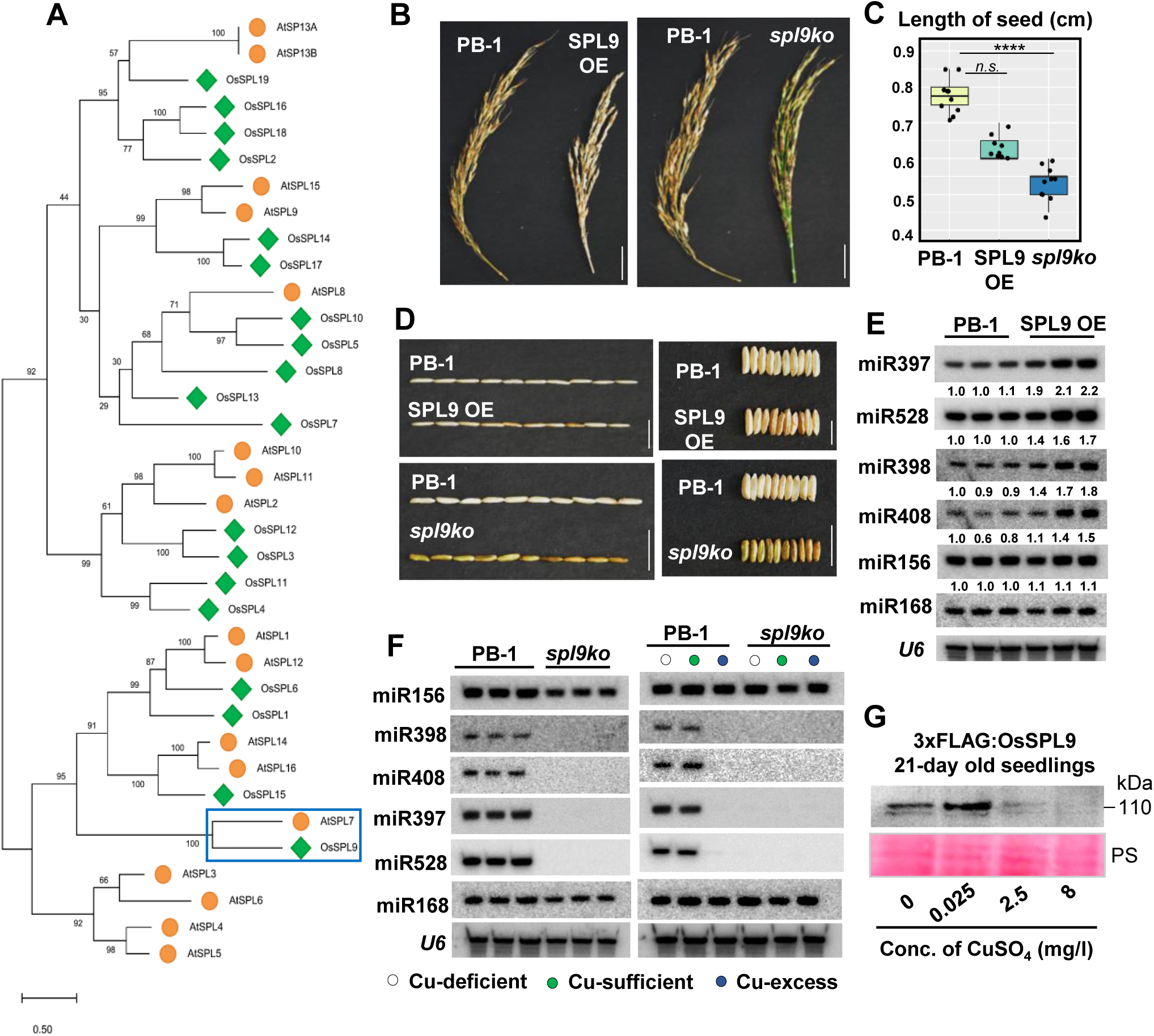
SPL9 regulates domestication associated Cu-miRNAs in rice. (A) Phylogenetic analysis of 16 AtSPLs and 19 OsSPLs indicated by orange circles and green inverted squares, respectively. AtSPL7 and OsSPL9 are highlighted. (B) Panicle size of SPL9 transgenic lines in PB-1 background. Size bar is 5 cm. (C) Box plot showing the seed length difference. Two-tailed Student’s t-test was used for statistical comparison. (*) p-value < 0.05, (ns) non-significant. (D) Seed phenotypes of SPL9 transgenic lines. Size bar is 5 cm. (E) Northern analysis for miRNAs in SPL9 OE lines. (F) miRNAs levels in *spl9ko* (left panel) and under varying Cu concentrations (indicated by coloured dots) (right panel). In E and F, *U6*, miR168 and miR156 served as controls. (G) Western blotting for OsSPL9 upon varying Cu concentrations. Ponceau (PS) served as loading control.

To study further the role of OsSPL9 in cultivated *indica* rice, we generated overexpression (OE) and knockout (ko) lines of OsSPL9. SPL9 constitutively expressed under *Zm:Ubi* promoter was tagged with 3xFLAG in the N-terminal (Supplementary Figure 3A) and the transcript/protein levels were validated (Supplementary Figure 3B-D). We obtained 8 independent transgenic lines upon transforming scutella derived PB-1 calli (Supplementary Table 4). Interestingly, the SPL9 OE plants showed more number of tillers (Supplementary Figure 3E), as well as reduced yield mainly due to reduced panicle size and abnormal grain filling. They also exhibited reduced seed length and width (Figure 3B-D).

We also generated *spl9ko* lines using a gRNA targeting the first exon (Supplementary Figure 3G) and obtained 5 independent ko lines (Supplementary Table 4), among which altered editing of SPL9 led to reduced levels of its transcript (Supplementary Figure 3H). The CRISPR-Cas9 mediated ko of SPL9 TF rendered a 2*nt*-deletion resulting in a truncated protein of 118 amino acids (Supplementary Figure 3I). The *spl9ko* plants were largely indistinguishable in vegetative growth when compared to WT (Supplementary Figure 3E), however, they exhibited compromised yield phenotypes such as reduced panicle size (Figure 3B). Seeds of *spl9ko* were also smaller and had abnormal shape (Figure 3C, D). These results indicate that mis-expression of SPL9 leads to alterations in yield related phenotypes such as seed length (Figure 3C), indicating the importance of optimal levels of SPL9 required for normal growth and development.

In order to understand if SPL9 is indeed upstream of Cu miRNAs, we performed northern blot analysis in these mis-expression lines and observed that all 4 miRNAs, but not other miRNAs, are upregulated at-least 2-fold in SPL9 OE lines (Figure 3E). The miRNA accumulation was also conserved across the vegetative leaf stages of the SPL9 OE lines (Supplementary Figure 3J). Most importantly, levels of these 4 miRNAs were undetectable in *spl9ko* in northern blot analysis (Figure 3F), together indicating that OsSPL9 is the upstream regulator of these miRNAs. Since we observed differential Cu accumulation between wild and cultivated lines, we explored if SPL9 mediated regulation of miRNAs gets altered under different levels of Cu. In wild-type PB-1, Cu-deficient conditions did not alter levels of these miRNAs, while Cu-excess conditions abolished the expression of all these miRNAs (Figure 3F). *spl9ko* lines did not express these miRNAs in both deficient and excess conditions, indicating that SPL9 is the sole regulator of these miRNAs under varying Cu conditions (Figure 3F). However, the SPL9 transcripts did not accumulate differently upon varying Cu concentrations in PB-1 (Supplementary Figure 3F). Interestingly, the high Cu led to reduced protein levels of SPL9 (Figure 3G) in agreement with a previous study (Yao et al. 2022), and this might be why miRNA expression was abolished under high Cu conditions. All these results conclusively indicate that SPL9 regulates the expression of Cu miRNAs involved in Cu-responses. OsSPL9 likely played a major role in establishing domestication-associated yield phenotypes by regulating the targets of these Cu-associated miRNAs.

### SPL9 binds to Cu-associated gene promoters enriched with the conserved GTAC motifs

Although SPL family of TFs in *Arabidopsis* are known to bind to GTAC containing motifs, global profile of regions bound by OsSPL9 or its other monocot homologs is unknown. Although it was shown that OsSPL9 can bind to miR528 promoter in ChIP-qPCR (Yao et al. 2019), it is unknown if it can also bind to promoters of other Cu-associated miRNAs and Cu-associated genes. Since SPL9 regulated the expression of miRNAs, we profiled SPL9 bound regions in wild type and under Cu-stressed conditions.

ChIP-seq was performed using seedling tissues using Illumina platform and we obtained more than 20 million reads mapping to IRGSP-1.0. We obtained more than 8000 peaks enriched with SPL9 binding. Among these peaks, most were located in intergenic regions, especially among promoters (63.5%), while distal intergenic regions (34.92%) peaks were also found in introns and exons (Figure 4A and B).Preferred binding to genic promoters by OsSPL9 is in agreement with similar observations in *Arabidopsis* (Schulten *et al*., 2022). The SPL9 bound peaks upon varying copper concentrations also showed intersection with genes (Figure 4C). The consensus motifs across the peaks were analysed and these had multiple GTAC motifs as expected, usually in the distal intergenic regions. Cu-sufficient conditions also showed wider stretches of GTAC motifs in SPL9-bound genomic regions (Figure 4D). Very interestingly, several Cu-associated genes, as well as several stress associated genes had SPL9 peaks in their promoter as well as in distal regions (Figure 4E). SPL9 appears to be regulated by itself as it bound to GTAC motifs in its promoter (Supplementary Figure 4).

**Figure 4.**
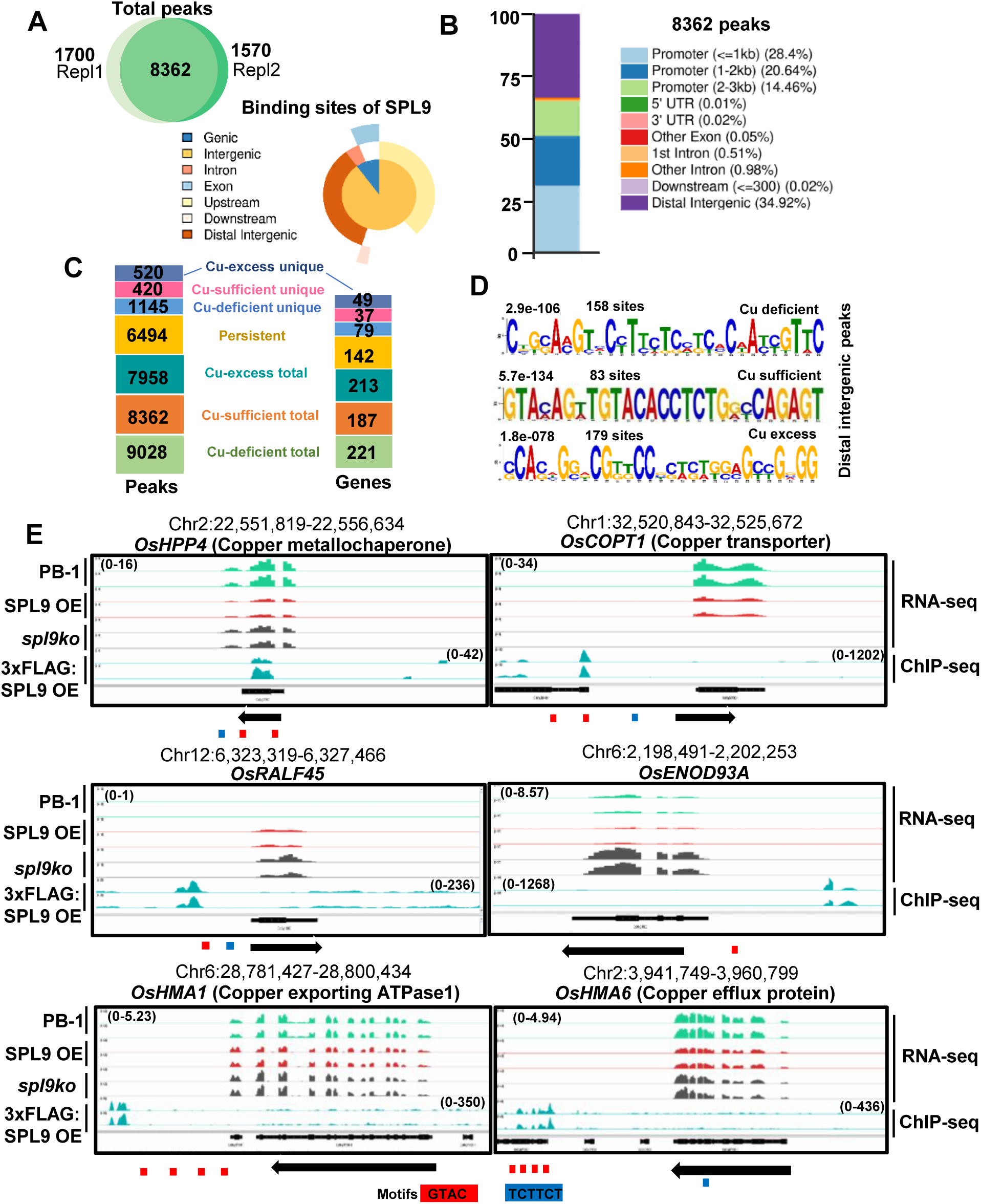
SPL9 TF binds to the consensus GTAC motifs near the protein-coding genes. (A) Venn diagram showing overlap of SPL9 peaks across replicates. The intersected peaks are represented denoting the occupancy under copper sufficient conditions. (B) Stacked barplot showing the percentage representation of the 8362 peaks across different genomic regions. (C) The genes corresponding to each peak is represented in the stacked bar diagram. (D) Representation of motifs across distal intergenic regions in Cu-deficient, Cu-sufficient and Cu-excess conditions. The e-value and the number of sites is mentioned. (E) IGV screenshots showing the occupancy of SPL9 across promoters and distal regions of several Cu-associated genes. The arrows indicate the orientation of the gene and the panels indicate RNA-seq (top) and ChIP-seq (bottom). Red blocks - GTAC motifs, blue blocks - TCTTCT motifs. The track data ranges are mentioned in parentheses.

Since miRNA and Cu-associated gene expressions were altered during stress, we hypothesized that SPL9 levels or binding preferences might get altered during Cu stress. SPL9 preferred peaks under varying concentrations of Cu were identified using ChIP-seq analysis and common as well as stress-specific peaks were intersected. We identified 9028 SPL9-bound peaks upon Cu-deficiency, 8362 peaks upon Cu-sufficiency and 7958 peaks upon Cu-excess conditions. Around 1145 unique peaks overlapped with 79 genes under Cu-deficient conditions. Similarly, around 420 unique peaks overlapped with 37 genes upon Cu-sufficient conditions and 520 unique peaks overlapped with 49 genes in Cu-excess conditions (Figure 5A). Although the peak width and intensity increased upon excess Cu conditions, number of SPL9-bound regions reduced marginally (Figure 5B and C). Among these peaks, 6494 peaks were persistent and unaffected by Cu stress and these overlapped with 142 genes (Figure 5D).

**Figure 5.**
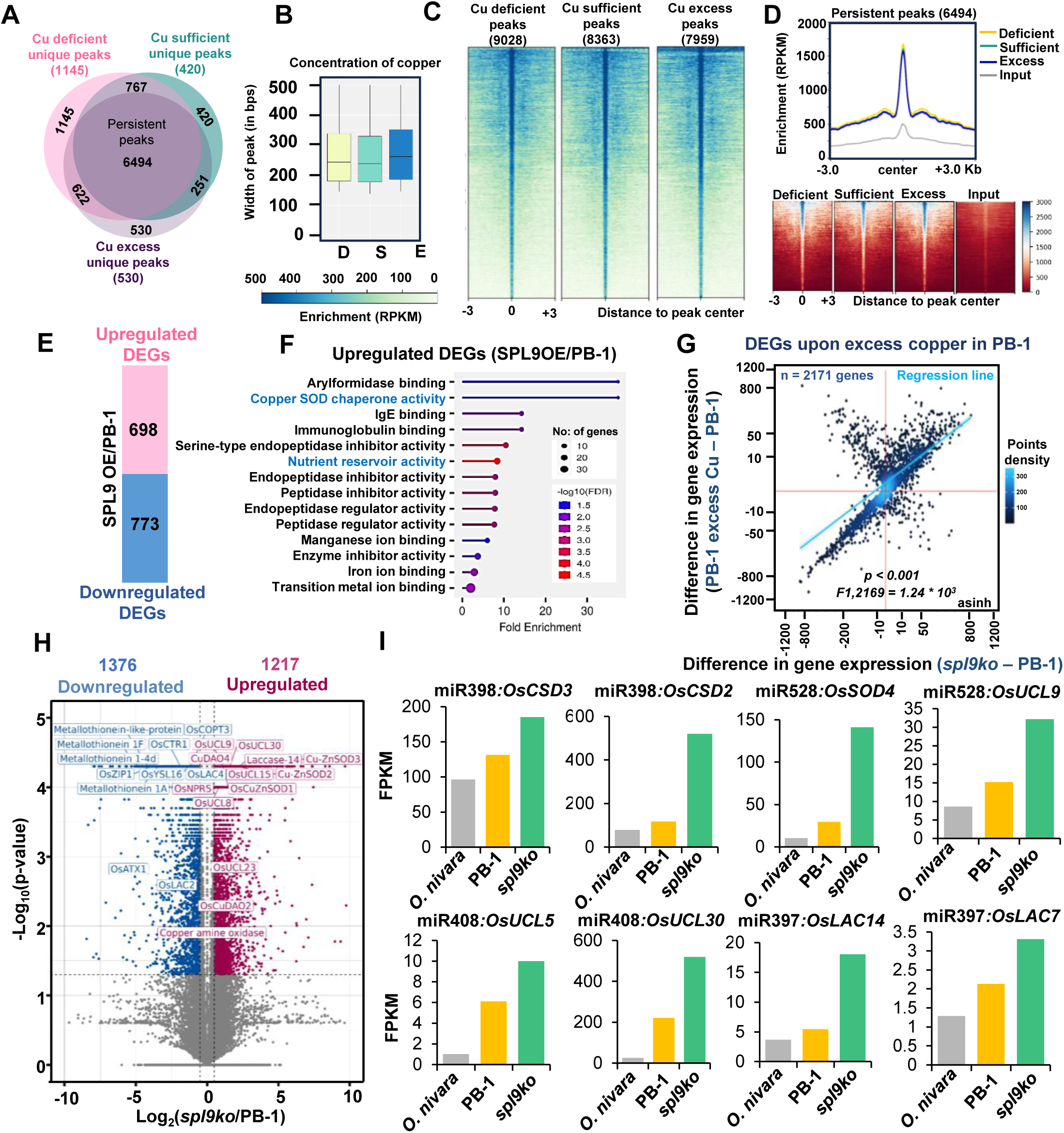
SPL9 binding to regulatory regions correlates with Cu-associated DEGs. (A) Overlap of ChIP-seq peaks under different Cu concentrations in 21-day old seedlings. (B) Box plot showing peak width across different Cu concentrations. (C) Metaplot showing total peaks and their distribution. (D) Metaplot showing persistent peaks (6494) and their enrichment. (E) Upregulated and downregulated DEGs in SPL9 OE lines with log_2_ fold change >1 and <1, respectively. (F) GO enrichment categories among the upregulated genes in SPL9 OE. FDR-False discovery rate. (G) Density scatter plot showing difference in expression of Cu-excess DEGs in *spl9ko*. Pearson correlation coefficient (R) and *p*-values are mentioned. Point density is shown in colour gradient. (H) Volcano plot showing DEGs in *spl9ko* seedlings. Blue dots-downregulated genes, red dot-upregulated genes. (I) Expression of miRNA targets in *O. nivara*, PB-1 and *spl9ko* based on cuffnorm FPKM values.

### OsSPL9 regulates expression of Cu-associated miRNAs and protein coding genes

To further understand the fundamental role of SPL9 TF in regulating gene expression under homeostatic conditions, we generated RNA-seq datasets (Supplementary Table 1) from both *spl9ko* and SPL9 OE lines. In SPL9 OE, 698 genes were upregulated and 773 genes were downregulated (Figure 5E). The upregulated genes belong to categories involved in Cu-chaperone associated superoxide activity, nutrient reservoir activity, enzyme inhibitor activity etc. and some of these were downregulated in *spl9ko* as expected (Figure 5F). Since OsSPL9 has specific roles in Cu-deficiency responses (Garcia-Molina et al. 2014; Wang et al. 2024b), we analysed the transcriptomic responses upon low Cu and excess Cu conditions in cultivated rice. The DEGs obtained in both the categories were correlated with the transcriptomic responses of *spl9ko*. Interestingly, we observed that *spl9ko* mimics an excess Cu transcriptome (Figure 5G).

The *spl9ko* lines showed 1217 upregulated and 1376 downregulated genes, the latter set of DEGs included copper transporter, COPT3 and several metallothionein proteins (Figure 5H). This is in agreement with observations in *Arabidopsis* SPL7 where the protein activated the expression of Cu-transporters such as *COPT1*, *COPT2* (Yamasaki et al. 2009). Increased Cu-miRNA target expression in *spl9ko* lines was also observed. The miRNA targets such as *CuZnSOD2*, *CuZnSOD3*, *SOD4*, *Uclacyanin 8*, *Uclacyanin 9*, *Uclacyanin 15, Uclacyanin 23*, *Uclacyanin 30*, *Laccase 14* which were upregulated several folds in *spl9ko* (Figure 5H). Among these, most of them showed reduced expression in *O. nivara*, while their levels were upregulated in *spl9ko,* when compared to PB-1 (Figure 5I). All these results conclusively indicate the importance of Cu-dependent processes mediated by OsSPL9 in homeostasis and under varying Cu-levels.

We further overlapped the upregulated and downregulated genes of *spl9ko* with SPL9 TF bound 8362 peaks identified in Cu-sufficient (optimal Cu) conditions. Upon overlapping *spl9ko* upregulated genes with SPL9 peaks, we obtained 57 genes corresponding to 158 peaks (Supplementary Figure 5A), all showing SPL9 binding in the distal regions of the genes (Supplementary Figure 5B, C). The 57 upregulated genes in *spl9ko* were also bound by SPL9, they were probably negatively regulated, as these genes failed to respond to low Cu conditions in PB-1 and in *spl9ko* (Supplementary Figure 5D). These genes were associated with defense response and protein desumoylation (Supplementary Figure 5E), and they were also directly or indirectly Cu-associated. The SPL9-bound upregulated genes included genes such as Iron-sulfur cluster protein, *Uclacyanin 23* etc (Supplementary Figure 5F) and they showed peak occupancy in the TES regions of coding sequences.

Upon overlapping *spl9ko* downregulated genes with SPL9 peaks, we obtained 66 genes corresponding to 208 peaks (Supplementary Figure 6A), all showing a higher SPL9 binding in the distal regions of the genes (Supplementary Figure 6B, C). In low Cu conditions, these 66 genes remained largely downregulated in *spl9ko* in comparison to PB-1. However, under excess Cu treatment, these 66 genes were predominantly downregulated in both *spl9ko* and PB-1. This conclusively indicated that these 66 genes are positively regulated at the transcriptional level by OsSPL9 in optimal Cu-conditions, and might have important roles to play under varying concentrations of Cu (Supplementary Figure 6D). In agreement with this prediction, gene ontology analysis revealed that these SPL9-bound downregulated genes (66) in *spl9ko* were associated with suberin biosynthesis, phenylpropanoid biosynthesis, fatty acyl biosynthetic process, etc. (Supplementary Figure 6E), and genes of these categories were among the DEGs upregulated in *O. nivara* when compared to PB-1 (Supplementary Figure 1B). It is important to note that these categories of genes are mediators of largely Cu-associated processes. Genes such as Heavy metal associated protein 2 (*HMA2*) and Pollen fertility restorer 2 were downregulated and they had SPL9 bound peaks close to TSS and TES (Supplementary Figure 6F).

### Cu-associated miRNAs form a feedback loop

Several studies have shown that stability of the miRNAs is regulated by their targets, association with Argonaute proteins or their modifications(Xie et al. 2003), indicating the presence of a feedback loop. A number of targets associated with Cu-binding and/or associated genes are regulated by the miRNAs downstream of SPL9.

However, the role of individual miRNAs or their targets in regulating the Cu-responses remained unclear. It is also intriguing that Cu, an element, the accumulation of which might vary between two nearby sites was central to these regulatory modules. We hypothesized that regulatory roles played by individual Cu-miRNA, its target(s) and Cu-availability/accumulation might be strongly linked in such a way that small perturbations do not lead to drastic growth penalty. In order to understand the crosstalk by SPL9-mediated regulation of miRNAs and its targets, we generated individual knockouts of these domestication-associated miRNAs in cultivated rice background (Supplementary Figure 7). The knockout of miR397 resulted in a 4 *nt* deletion in the stem loop of the precursor miRNA, perturbing the formation of mature miR397 (Supplementary Figure 7A). *miR528ko* mutant harbours a single nucleotide insertion (Supplementary Figure 7B) and the knockout of miR408 using CRISPR led to 8 *nt* deletion in the stem-loop region of the miRNA (Supplementary Figure 7C). The mature miR408 with this mutation will be only 18 *nt* long when compared to 21 *nt* in WT. All these knockout plants showed severe growth phenotypes when compared to the WT even in seedling stage, indicating the role of these miRNAs and their targets in early development (Figure 6A-D).

**Figure 6.**
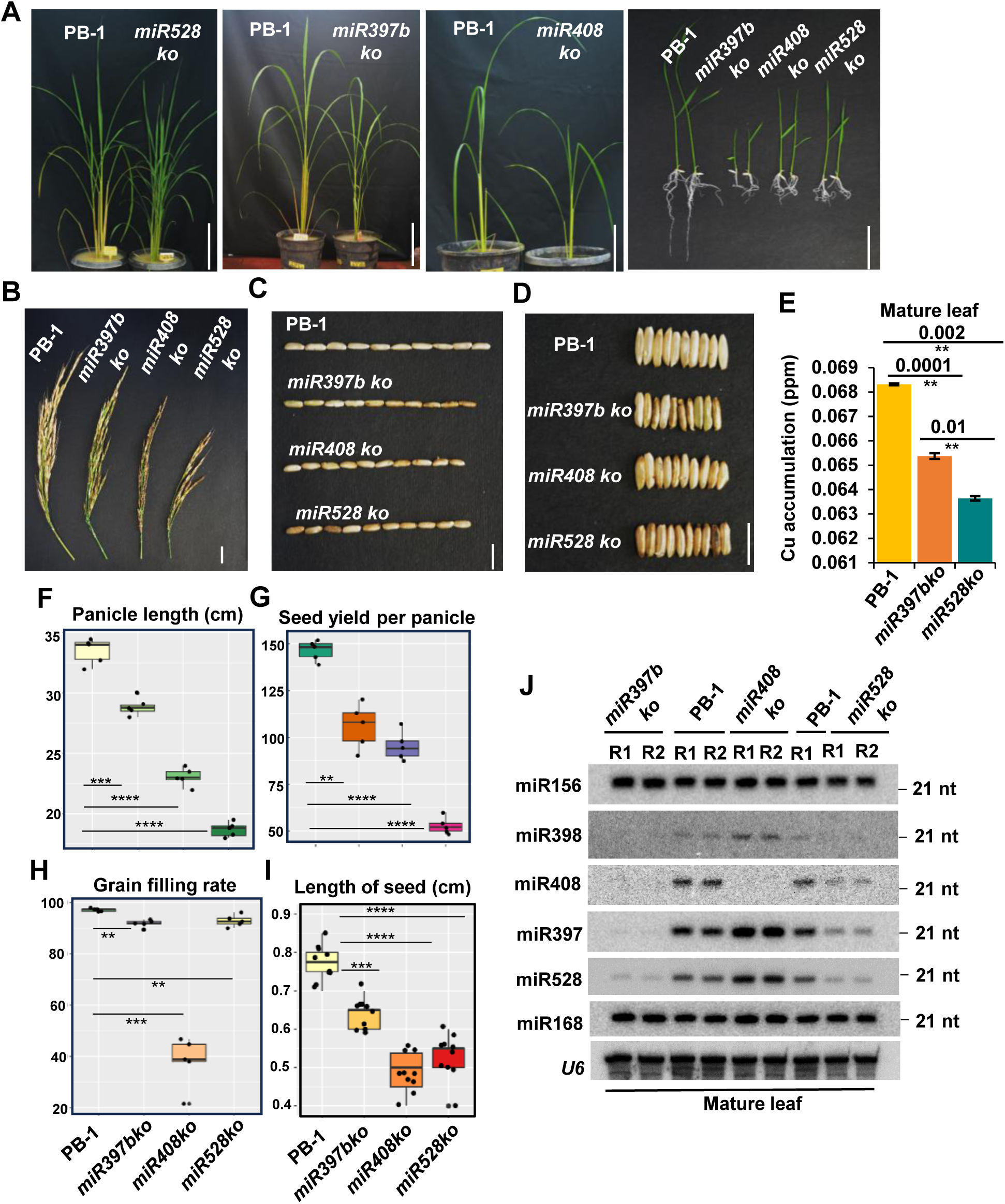
Cu-responsive miRNA mutants show compromised yield and suppression of other Cu-miRNAs. (A) miR397b, miR528, and miR408 ko lines show developmental phenotypes. (B) Phenotypes of panicles in various miRNA mutants. (C) Phenotypes of seed length in various miRNA mutants. (D) Seed width in various miRNA mutants. (E) Accumulation of Cu in miRNA ko lines in leaf tissues. (F) Box plot showing panicle length among miRNA mutants. (G) Box plot showing seed yield per panicle among mutants. (H) Box plot showing grain-filling rate among miRNA mutants. (I) Box plot showing seed length in miRNA mutants. (F-I) Two-tailed Student’s t-test was used for statistical comparison. (*) *p*-value < 0.05, (*ns*) non-significant. (J) Northern blotting showing miRNA accumulation in miRNA mutants. *U6*, miR168 and miR156 served as loading control for miRNAs.

Estimation of the Cu accumulation revealed that in the mature leaves of miRNA knockout lines of miR528 and miR397, there was reduction in Cu accumulation (Figure 6E) similar to SPL9 perturbed lines. This suggested a strong feedback control between individual miRNAs and their regulator OsSPL9 as it indicates that the SPL9-regulated phenotypes are dependent on SPL9-regulated miRNAs. On a similar note, the panicle length, seed length and width were reduced in each of these knockout lines, suggesting that individual miRNA knockouts phenocopied *spl9ko* lines (Figure 6F-I).

The perturbation of individual miRNAs led to changes similar to their upstream regulator SPL9, it is likely that each Cu-associated miRNA module functions in a feedback regulation with SPL9 and in turn, other SPL9-dependent miRNAs. This probed us to test the accumulation of other SPL9-dependent, domestication-associated miRNAs in individual miRNA knockout lines. Very interestingly, we observed that the individual miRNA knockouts of miR397 and miR528 also led to reduction in all the other domestication-associated miRNAs (Figure 6J). For example, the *miR397bko* not just led to reduction in the levels of miR397b but also the other Cu-associated miRNAs such as miR528, miR408 and miR398. Such a relationship was observed partially or fully with other Cu-associated miRNA knockouts. For example, *miR408ko* did not alter the levels of other Cu-associated miRNAs but only that of miR408. However, similar to *miR397ko*, *miR528ko* accumulated reduced levels of miR528 and also the other Cu-associated miRNAs. This is indicative of a strong self-regulatory module where the regulation of miRNAs is not only dependent on each other but also on the targets or the set of Cu-genes regulated by them. Since such feedback is not reported so far, it will be interesting to identify how upstream regulators of Cu-miRNAs influence miRNA biogenesis machinery. Expression levels of genes involved in miRNA biogenesis remain unperturbed under varying Cu-levels (Supplementary figure 7D). In *hen-1* mutants, the transcripts of Cu-binding proteins such as *LAC5* and *LAC8* accumulated independent of copper supply and to higher levels compared to wild type (Abdel-Ghany and Pilon 2008). It is possible that stability of proteins, especially those involved in miRNA accumulation, might be under the control of proteases as hypothesized previously for SPL7 (Garcia Molina et al., 2014).

### Perturbation of Cu levels in wild and cultivated lines led to yield compromised phenotypes

In order to understand the role of Cu in mediating SPL9 and domestication-associated miRNAs, we subjected wild and cultivated rice lines to different concentrations of Cu (Figure 7A). Levels of miRNAs across wild and cultivated lines in Cu-deficient, sufficient and excess conditions were assayed using northern analysis. Under Cu-deficient conditions, the levels of domestication-associated miRNAs remained unchanged in *O. nivara*, while their expression was induced in PB-1. However, upon Cu-excess conditions, miRNA levels were reduced in both *O. nivara* and PB-1 (Figure 7B). Similarly, SPL9 OE lines mimicked *O. nivara* in that the levels of Cu-miRNAs were unaffected in Cu-deficient conditions, whereas, Cu-excess led to reduction in miRNAs. However, partial retention of miRNA levels of miR528 and miR397 were observed in SPL OE lines only upon high Cu, indicating that the levels of SPL9 protein dictates the miRNA levels. It is possible that miRNA targets including those required for Cu-binding proteins are already downregulated in *O. nivara* due to higher accumulation of miRNAs.

**Figure 7.**
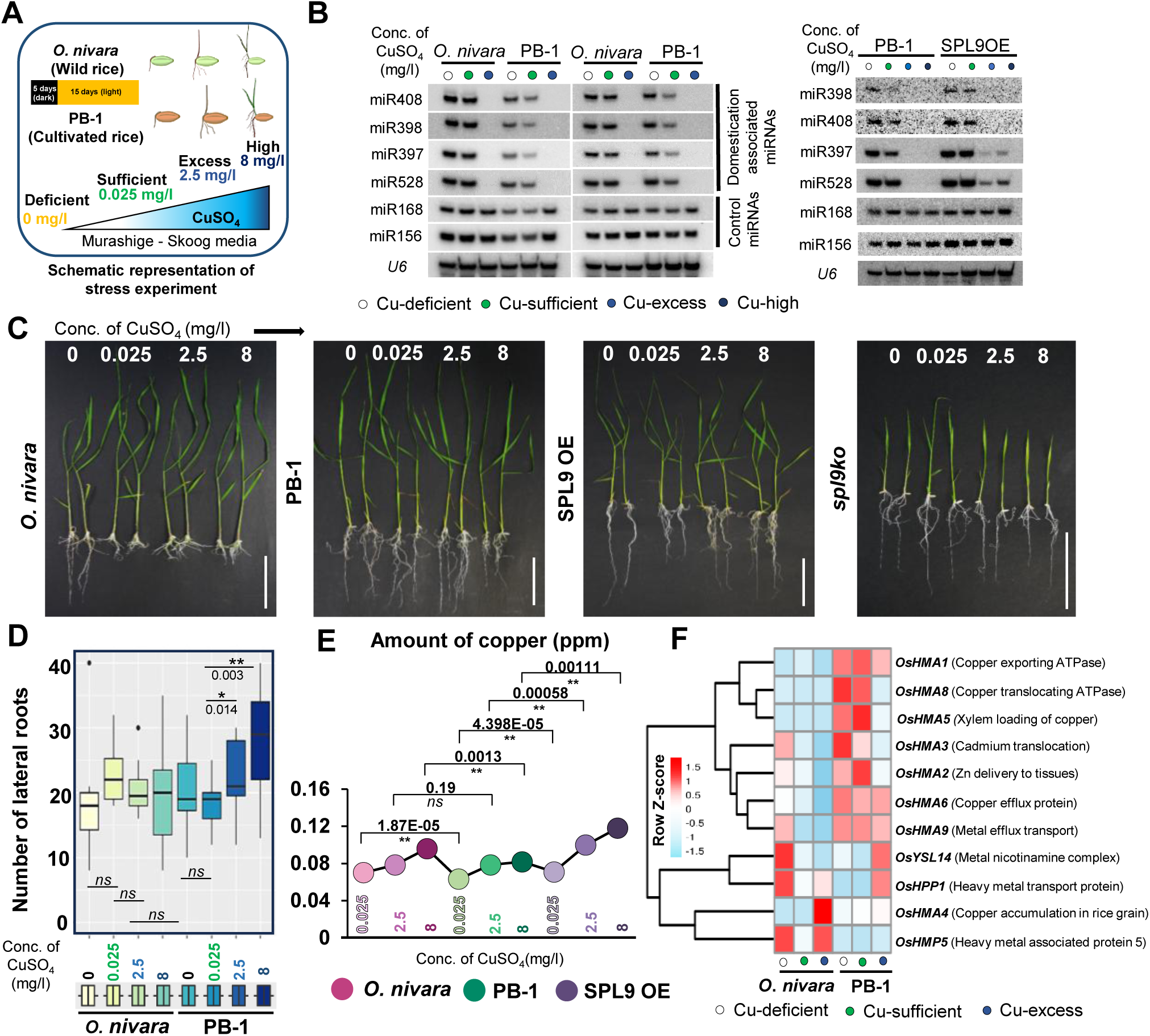
Cu-stress induces differential yield-related phenotypes between *O. nivara* and PB-1. (A) Schematic representation of Cu stress in wild and cultivated lines. (B) Northern analysis showing abundance of miRNAs. miR156, *U6* and miR168 served as loading controls. (C) Phenotypes of wild and cultivated rice post-exposure to Cu stress. (D) Box plots showing the lateral root length upon Cu stress in *O. nivara* and PB-1. (E) Cu accumulation in wild and cultivated rice upon Cu stress. X-axis indicates concentration of copper across samples, Y-axis indicates amount of copper in parts per million (ppm). (D-E) Two-tailed Student’s *t*-test was used for statistical comparison. (*) *p*-value < 0.05, (*ns*) non-significant. (F) Heatmap showing the expression of HMA genes. The dots indicate increasing Cu concentrations.

**Figure 8.**
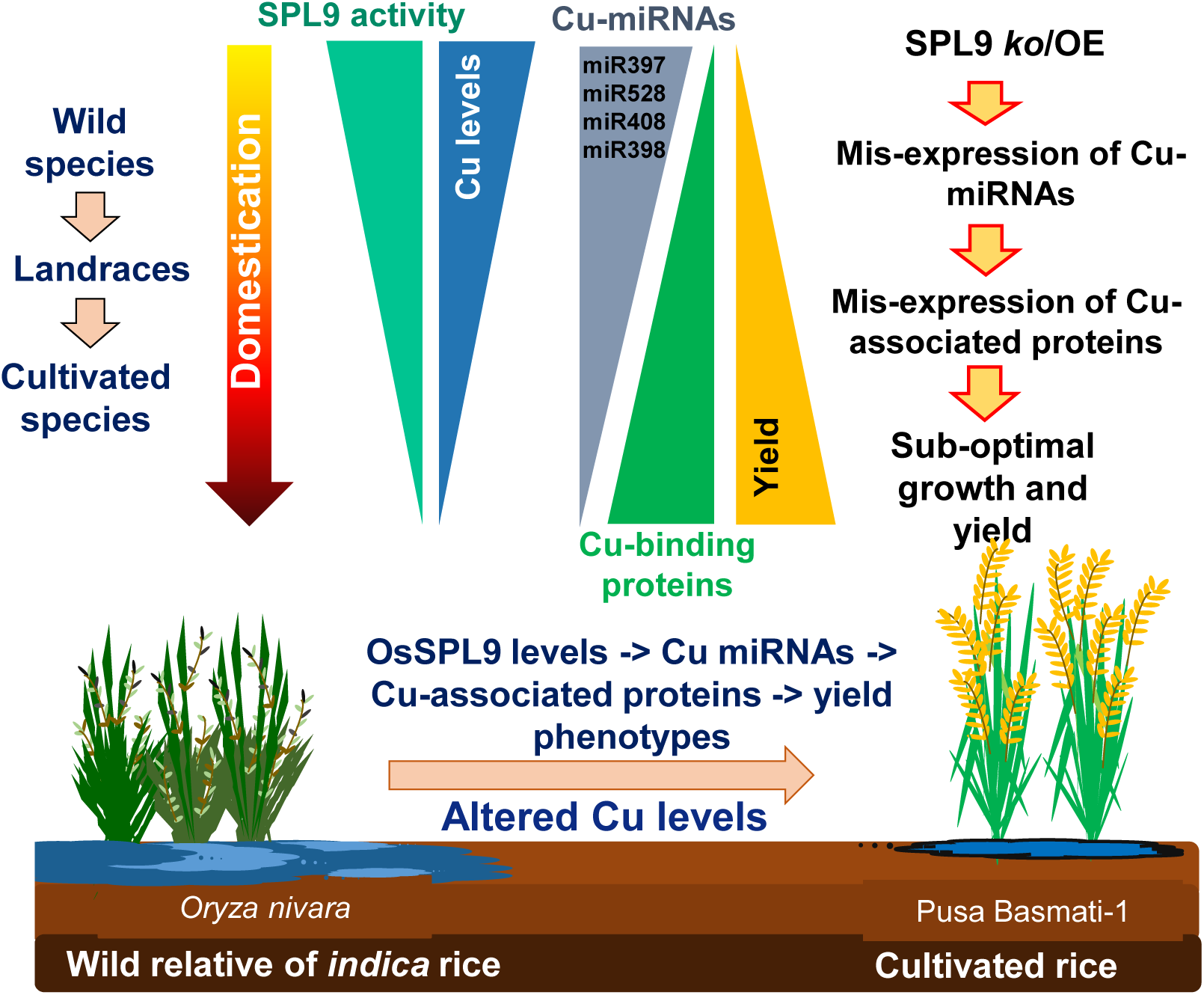
Proposed model for Cu-mediated regulation of domestication-associated phenotypes in *indica* rice. Depicted is the schematic representation of the contribution of Cu levels, SPL9, Cu-miRNAs and Cu-associated proteins in regulating domestication associated phenotypes in *indica* rice.

However, the primary root length did not change drastically in wild rice compared to the cultivated rice under Cu-excess conditions, while root length increased under deficient conditions in *O. nivara*, SPL9 OE but not in PB-1 and *spl9ko* (Figure 7C). The cultivated rice exhibited robust lateral root growth (Figure 7D) upon increasing Cu concentrations in agreement with earlier studies (Lequeux et al. 2010). In the wild rice, the lateral root growth was unaffected upon increasing Cu concentrations. All these observations indicate that seedling growth is differentially regulated between wild and cultivated rice lines under varying Cu-levels. *O. nivara* and SPL9 OE had 421 DEGs in common (Supplementary Figure 8A), and these were majorly associated with oxidoreductase activity (Supplementary Figure 8B). Correlation analysis also indicated that a major category of genes contributing to the transcriptomic response upon excess Cu is conserved across SPL9 OE and the wild rice, *O. nivara* (Supplementary Figure 8C). Together, these indicate the important role of OsSPL9 in modulating miRNAs and copper-responsive genes in a Cu-dependent manner.

To identify the nature of these conserved genes that overlap between *O. nivara* and SPL9 OE, the upregulated and downregulated genes of both were considered. There were 185 genes downregulated in *O. nivara* and SPL9 OE in comparison to PB-1 (Supplementary Figure 8D). Several ion channel membrane transporters were in this category. Among the 161 shared upregulated DEGs between *O. nivara* and SPL9 OE (Supplementary Figure 8E), there were genes coding for regulators having oxidoreductase activity. Very few genes among the shared DEGs showed opposing pattern of gene expression between *O. nivara* and SPL9 OE (Supplementary Figure 8F, G).

In order to correlate specific phenotypes under the control of Cu-dependent genes, we tested the accumulation of SOD, a major regulator of oxidative stress and signalling, upon Cu-stress. Expression of Cu-SOD1, a target of miR398 and Cu-SOD4, a target of miR528, were highly expressed in PB-1 as expected due to low Cu-miRNAs. As expected, levels of H_2_O_2_ as measured using 3,3’-diaminobenzidine (DAB) staining, showed increased accumulation of brown precipitates in *O. nivara* matching reduced SOD levels (Supplementary Figure 9A). Expression of these SODs were induced in PB-1, but not in *O. nivara* under Cu-excess conditions (Supplementary Figure 9B). Correspondingly, the levels of anthocyanin that is routinely associated with oxidative stress tolerance among plants, accumulated to higher levels in *O. nivara* when compared to PB-1. As expected, expression of anthocyanidin and proanthocyanidin synthase genes were highly expressed in *O. nivara* compared to PB-1 (Supplementary Figure 9C). Similar observations were made when the plants were subjected to Cu-stress (Supplementary Figure 9D).

Since untreated samples of wild and cultivated lines exhibited differences in Cu-levels, we explored if wild and cultivated rice lines accumulate Cu differentially under Cu-stress. Wild and cultivated lines across increasing Cu concentrations showed an increased accumulation of Cu, whereas the wild rice, *O. nivara* accumulated Cu somewhat similar to SPL9 OE lines (Figure 7E). The levels of Cu accumulation in PB-1 were reduced when compared to that of the wild rice *O. nivara* and SPL9 OE lines. Transcriptomic analysis also indicated that several HMA transporters such as HMA1, HMA6, HMA8, genes that are Cu-exporting/translocating ATPases, are highly expressed in cultivated rice when compared to *O. nivara* (Figure 7F). All these results indicate levels of Cu as a major determinant of differences in Cu-miRNA expression, expression of Cu-associated genes and targets of miRNAs, and Cu-dependent phenotypes. These suggest unappreciated roles of Cu in modulating *indica* rice domestication-associated phenotypes.

## Discussion

Cereal crops grown in agricultural ecosystems are genetically distinct from wild relatives and are subjected to varied water and fertiliser supplies. Ability to acquire and utilize the minimal requirements for plant growth, must have contributed to the process of domestication in these crops, to enable them to have high photosynthetic rate, broader leaves, and increased growth rate (Chamberlain-Irwin and Hufford 2022; Huang et al. 2022a; Alam and Purugganan 2024). Increased growth might have demanded more nitrogen and phosphorus uptake and utilization (Krouk and Kiba 2020), when compared to other elements such as Cu and micronutrients that are part of defense proteins and promoters of secondary metabolites (Edmondson and Thimann 1950). Uptake and accumulation of several of these elements, including metal ions, were altered in many agronomic crops during domestication (Muñoz et al. 2016). Studies have suggested that genetic as well as epigenetic factors alter the histone marks at the level of transport, metabolism as well as signalling for macronutrients such as nitrogen and phosphorus (Li et al. 2021a). Cu being a micronutrient, an essential co-factor for a wide range of cellular enzymes and proteins, has importance in several key processes. Cu-proteins are functionally redundant to existing iron enzymes and have evolved much later upon oxygenation, indicating their possible redundant but equally important roles in the metalloproteomes (Schulten et al. 2019). It is therefore not surprising that Cu build-up and its utilization and incorporation appear to be a major determinant for traits somewhat similar to the well-established macronutrient nitrogen (Sun et al. 2014; Chao and Lin 2015; Brasier et al. 2020; Lei et al. 2021; Garcia et al. 2022; Huang et al. 2022b; Hu et al. 2023a). Even though Zn, Fe and Mn associated metalloproteins exist, there is no clear association of them being key regulators of domestication-associated traits (Kumar et al. 2021). It is evident that Cu might have a bigger unappreciated role in plant growth and development than previously envisioned. Accumulation of high Cu in wild relatives is in agreement with their ability to have stronger disease resistance traits (Gasparini et al. 2021). Further studies are required to understand the nature of its accumulation in tissues, subcellular localization, stored and utilised form of Cu, as well as some understanding of Cu cycle and its homeostasis within wild and cultivated rice plants.

The TFs function by binding to contact sites in the enormous stretch of DNA to promote the recruitment of other factors such as co-activators/repressors to regulate the gene expression (Franco-Zorrilla et al. 2014; Lis and Walther 2016; Strader et al. 2022). SPL family of TFs are well known in rice to modulate yield associated traits, for example, OsSPL14 is important for regulating plant architecture (Jiao et al. 2010; Luo et al. 2012; Guo et al. 2021), and SPL12 and miR529-regulated SPLs such as SPL2, 16, 17, 18 are essential for maintaining grain size and yield (Qin et al. 2020; Yan et al. 2021). Maize SPL homolog regulates kernel row number (Wei et al. 2018). Among dicots too, SPLs are well-known regulators of growth (Ferreira e Silva et al. 2014). Since Cu metabolism in rice is connected closely with yield-related traits and miRNAs form feedback to regulate them to a very refined layer through OsSPL9, it is clear that SPLs have the potential to modulate Cu-mediated processes. Similar to other SPL members (Wang et al. 2009; Yamasaki et al. 2009; Araki et al. 2018; Gou et al. 2019), OsSPL9 also preferred GTAC motifs. We have also observed that there are additional sequence motifs where SPL binding can be seen and it will be interesting to further explore if these additional motifs mediate specific functions. In this study, the perturbation of OsSPL9 or even altering amount of Cu available, mimicked the accumulation of miRNAs/target levels in wild or cultivated rice. This conclusively indicates that Cu-dependent SPL9 mediated regulation is central to gene expression differences between wild and cultivated rice lines.

Homolog of OsSPL9 in *Arabidopsis*, SPL7, is known to interact with another TF named HY5, thereby integrating the light and Cu cues. The pathway is also regulated by miR408 such that it reallocates more Cu to chloroplasts (Zhang et al. 2014). The sequence specific DNA-binding activity of the OsSPL9 TF indicates the ability to regulate the miRNA genes as well as several Cu proteins/transporters in a synchronous manner. This required optimal levels of OsSPL9 as reported here, since Cu-stress altered binding to target sites. It will be interesting to study if OsSPL9 integrates other cues such as light and temperature to modulate binding to target sites. Some of the downstream targets of OsSPL9 seem to be working in this fashion. For example, Cu-associated miR528 targets RFI2 (Red and far-red insensitive 2), a RING-domain zinc finger protein that modulates light sensing (Yang et al. 2019). Natural variations in the miR528-SOD interaction has been documented in C3 and C4 photosynthetic monocot species (Han et al. 2024). Additionally, *Arabidopsis* miR408 encoded peptide miPEP408 accumulated differentially during light transitions (Kumar et al. 2024). miR398 is a heat-responsive miRNA that facilitates thermotolerance among diverse plants (Li et al. 2020). It is possible that through these regulatory loops, OsSPL9 regulation might be controlled by other environmental cues apart from natural variation, aging and light as reported previously (Yang et al. 2019).

Since OsSPL9 appears to be a major regulator of yield related traits, it is possible that its homologs in other cereals also regulate Cu-dependent yield-related processes. Although such studies are yet to be carried out in cereals other than rice, CRR1 gene, a homolog of OsSPL9 and AtSPL7 in *Chlamydomonas*, regulates Cu uptake and its utilization (Sommer et al. 2010). In moss, Cu-mediated transcriptional suppression is through a SBP transcription factor, PpSBP2 in a GTACT motif-dependent manner (Nagae et al. 2008). In maize, miR528 is known to regulate laccases and affect the lodging resistance (Sun et al. 2018), but the upstream players are unknown. In crops like banana, miR528-PPO module involved in fine-tuning ROS levels is under the regulation of a miR156 targeted SPL4 (Kong et al. 2025).

In this study, we observe feedback between individual miRNAs and OsSPL9 in a Cu-dependent manner. Several studies have pointed out feedback mechanisms where the miRNA machinery is regulated by a miRNA itself such as DCL1 and AGO1 negatively regulated by miR162 and miR168 respectively (Vaucheret et al. 2004). In the context of nutrient stress such as phosphate, the non-protein gene IPS1, acts as a target mimic to sequester miR399 leading to PHO2, its target upregulation (Franco-Zorrilla et al. 2007). Target directed miRNA decay which leads to miRNAs of distinct families getting degraded has been reported from metazoans (Hiers et al. 2024a, 2024b). In SPL9 mis-expressed lines, or in miRNA ko lines, levels of miRNAs other than Cu-miRNAs were unchanged. Among individual miRNA ko plants, absence of a miRNA might have triggered downregulation of other miRNAs in the pathway, probably by altering functions of a Cu-mediated process downstream of the core miRNA biogenesis machinery. It is also possible that levels of Cu incorporated by any of the miRNA targets in miRNA ko line, in specific cell types, might influence levels of Cu available for other Cu-associated proteins.

The miRNAs and TF responding to varying levels of Cu might have been a key for survival as Cu is an important micronutrient that serves as a cofactor for several enzymes and proteins (Burkhead et al. 2009). The evolution of such a system might be a result of the habitat in which these lines evolved and in soils enriched with specific metals. Several metabolic pathways and metabolites are triggered upon heavy metal stress in cultivated rice (Ahsan et al. 2007; Meier et al. 2012; Cao et al. 2023). However, *O. nivara* has higher accumulation of metabolites such as anthocyanins that can incorporate high amounts of heavy metals (Li et al. 2021b; Naing and Kim 2021; Kaur et al. 2023; Sun et al. 2023) to sequester them in vacuoles and less sensitive organelles/tissues. Over the course of domestication, the modern crop varieties have been prone to grow in nutrient-rich environments (Meyer et al. 2012; Hancock et al. 2025). It is likely that OsSPL9 played a crucial part in this selection and adaptability to agricultural systems.

## Materials and Methods

### Plant material

The plant materials used for the study were *Oryza nivara* and *Oryza sativa indica* PB-1 which were grown in a growth chamber 24 °C/70 % RH with a 16 h–8 h light– dark cycle. The plants grown in the growth chamber were further transferred to a greenhouse maintained at 28 °C with a natural day-night cycle. The other mutants in the study are mentioned in the Supplementary Table 4.

### Binary vector construction and *Agrobacterium* mobilization

For the generation of overexpression lines of SPL9, the CDS of the gene Chr 5: 19923167-19932333 (Os05g0408200) was amplified and cloned with an N-terminal 3xFLAG tag into a derivative of pCAMBIA1300 driven by a maize Ubiquitin1 promoter. The *spl9ko* mutant was generated using CRISPR-Cas9 mutagenesis and the gRNA was designed using CRISPR-PLANT (Minkenberg et al. 2019). A similar strategy was followed for the miRNA knockouts as well. The gRNA sequences and the reverse primers for all constructs are mentioned in the Supplementary Table 5. The amplicon harbouring the gRNA sequence was cloned and inserted to plasmid pRGEB32 (Xie and Yang 2013). Screening of mutants was performed by PCR and sequencing to identify the indel (insertion/deletion) in the targeted region.

The constructs were mobilized into Agrobacterium using the electroporation method. All the binary plasmids were mobilised into Agrobacterium tumefaciens strain LBA4404 harbouring extra-virulence plasmid pSB1. Embryogenic rice calli were infected with Agrobacterium strain containing binary plasmids using established methods (Hiei et al. 1994; Sridevi et al. 2003) and the calli were selected and regenerated on hygromycin containing medium.

### Stress experiments of the seedlings

About 21-day old seedlings grown on ½ MS media (without CuSO_4_) supplemented with increasing gradients of CuSO_4_ concentration (0 mg/l – Cu-deficient, 0.025 mg/l – Cu-sufficient, 2.5 mg/l – Cu-excess, and 8 mg/l – Cu-high) were used for experiments. The samples were collected post stress administration and snap frozen in liquid nitrogen for RNA extraction and other molecular analysis. For ChIP-seq analysis, the samples were crosslinked and processed. The phenotyping of the seedlings was performed by counting the root length, shoot length, number of lateral root hairs, total number of crown roots. The box plots were plotted using R-package ggplot2 and the statistical significance was computed using Student’s *t*-test.

### Cu estimation in seedlings

The rice seedlings grown in the stress media were collected at the end of stress regimen. The tissues were dried for a period of 3-5 days at 37 °C to remove moisture. At the end of drying period, the samples were finely ground to a powder using shredder. Around 200 mg of the ground powder was weighed and 10 ml of concentrated HNO_3_ was added. The tissue was digested overnight in a fume hood. Post overnight incubation, 1 ml of 4% H_2_O_2_ was added to the acid-digested tissue. Post digestion, the tissues were then put in an acid digestor to facilitate maximum extraction of the elements. The samples were filtered using a filter paper and collected. The samples were analysed along with Cu standards in an ICP-OES as described in (Yao et al. 2022).

### RNA-seq analysis

Total RNA was extracted from 21-day old whole seedlings. A total of 1 µg RNA was taken as input and RNA integrity was verified with Agilent Tapestation. RNA was enriched for poly-A fraction and library was prepared using NEBNext Ultra II Directional RNA Library Prep kit (E7765L) as per manufacturer’s instructions. Sequencing was performed in a paired-end fashion on Novaseq 6000 platform The resulting reads from the library were trimmed using Trimmomatic (Bolger et al. 2014) and mapped to the rice genome (IRGSP 1.0) using Hisat2 (Kim et al. 2015). The library statistics and the alignment percentage are mentioned in the Supplementary Table 1. The output files were later processed using Samtools (Cock et al. 2015) and Cufflinks package (Trapnell et al. 2012; Ghosh and Chan 2016). The DEG calling was performed using Cuffdiff and the upregulated and downregulated genes were classified based on log_2_Fold change cut-off as >1 and <-1. The p-value was taken as less than 0.05 for significant DEGs. For volcano plot, ggVolcano (Mullan et al. 2021) tool was used. For heatmap, pheatmap package in R was used. The scatter plots were made using ggplot2 after creating the matrix for the gene expression using bedtools multicov as mentioned in (Vivek et al. 2023). GO analysis was performed using Shiny GO v0.75 (Ge et al. 2020). The gene-IDs were used from RAPDB with a preference for biological processes with FDR cut-off of 0.05.

### sRNA northern hybridization

sRNA northern blots were performed based on the previously established protocols (Shivaprasad et al. 2012a; Tirumalai et al. 2020). Around 10 µg of TRIzol extracted total RNA from 21-day old seedling tissues was electrophoresed on a denaturing 15 % acrylamide gel. The RNA gel was blotted onto Hybond N+ membrane (GE Healthcare) and UV cross-linked. The hybridisation of the membrane was performed with the T4 PNK end-labeled oligonucleotides ([γ-^32^P]-ATP) in Ultrahyb buffer (Invitrogen). The blots were scanned using a Typhoon scanner (GE Healthcare). After the scanning, blots were stripped at 80 °C in stripping buffer and repeat hybridizations were done with subsequent probes.

### Phylogenetic analysis

Amino acid sequences in Arabidopsis and rice were downloaded from TAIR (https://www.arabidopsis.org/) and RAP-DB (https://rapdb.dna.affrc.go.jp/) respectively. A maximum likelihood phylogenetic tree was generated based on the alignment using MEGA v6.06 with bootstrap of (1000 repetitions).

### Chromatin immunoprecipitation-sequencing (ChIP-seq) and analysis

ChIP was performed as described earlier (Saleh et al. 2008; Song et al. 2016; Hari Sundar G et al. 2023). About 1-1.5 g of the 21-day old seedling tissue was taken and cross-linked with 1% formaldehyde with intermittent mixing and quenched with 0.125M glycine in 1X PBS. The tissues were finely ground in liquid nitrogen and the nuclei isolation was performed. The nuclei were lysed and sheared using Covaris ultrasonicator till the fragments were of 150-350 bp in size. The chromatin post shearing was incubated overnight with Anti-FLAG M2 Magnetic Beads (Sigma-M8823) at 4°C. The purified IP products were taken for library preparation using NEBNext Ultra II DNA library prep kit with sample purification beads (NEB E7103L) as per the manufacturer’s protocol. The libraries (with replicates) were sequenced on an Illumina Novaseq 6000 platform (Supplementary Table 3).

The datasets from the sequencing were used for cutadapt-based adapter trimming. The alignment was performed using Bowtie2 (Langmead and Salzberg 2012) and the alignment parameters were as follows: -v 1 -k 1 -y -a --best –strata. The PCR duplicates were removed and the alignment files were converted to coverage files and normalised with input samples, using deeptools (Ramírez et al. 2014, 2016). ComputeMatrix tool was used to compute the average signals over the annotated regions and plotprofile was used to generate the metaplots. The peak calling was performed using MACS2 with narrow peak calling option. The consistency across replicates was validated using BEDTools intersect. Intervene was used to obtain the intersections with the RNA-seq data as performed earlier (Harshith et al. 2024).

### sRNA sequencing and analysis

sRNA datasets were obtained from (Swetha et al. 2018). The size based fractionation and library preparation were performed as previously described (Tirumalai et al. 2019). The library statistics and alignment percentage are as mentioned in Supplementary table 2. These reads were further subjected to quality check using FastQC, adapter removal and size selection using UEA sRNA workbench (Stocks et al. 2012). The 21-22 nt size classes of the miRNA was categorized to IRGSP1.0 genome using Bowtie 2 using the following parameters: -v 1 -m 100 -y -a --best –strata. The miRNA abundance was profiled using miRProf tool (Stocks et al. 2018) and the differential expression of the reads were plotted. Only the mapped reads were used for further analyses, including differential expression analyses.

### Degradome analysis

The degradome library preparation and sequencing are as mentioned in (Swetha et al. 2022). In order to predict the potential miRNA targets, GSTAr script from CleaveLand was used. The degradome reads were aligned to all rice transcripts from MSU Rice Genome Annotation Project Version-7 using Bowtie with 2 mismatches. The calculation of fragment abundance of the reads and category for each position in every gene was performed. Later, the merging of the potential targeting information and read alignments from the genome was done to obtain the list of valid miRNA targets. The list of the potential targets had target cut site, category, and abundance. Category 0 is when more than 1 read aligns at the cut site and is the only position where maximum reads have aligned. Category 1 is when more than one read aligns and is also where the maximum number of reads align and is among the other equal maximum peaks. Category 2 is when more than one read aligns at the cut site and the depth is above the average depth (average of abundances at all positions that have at least one read) and below the maximum abundance. Category 3 is when more than one read aligns at the cut site and the depth is below or equal to the average depth (average of abundances at all positions that have at least one read). Category 4 is when only one read aligns at the cut position (Tirumalai et al. 2019).

### Western blotting

The mature leaves and seedling tissues were used for protein extraction and western blotting. The samples were finely ground using liquid nitrogen and extraction was performed using extraction buffer (0.05M Tris pH 8.0, 0.2M NaCl, 10% Glycerol, 1mM EDTA, 1% Triton-X-100, 1mM PMSF, PVP). Around 200 mg of the whole seedling or mature leaves were taken for the native extraction. The protein extract from the samples were electrophoresed stained using a CBBS to normalise the loading. Normalised amounts of proteins were electrophoresed and were transferred to nitrocellulose membrane. The membrane was hybridised with Anti-FLAG M2 monoclonal antibody (Sigma F1804) and detected using HRP-based chemiluminescent detection system.

### RT-qPCR

Around 1-1.5 µg of total RNA was used for cDNA synthesis and RT-qPCR. The qPCR reaction was set up using SYBR containing reaction mix (Solis Biodyne) and the samples were loaded as triplicates in a 96-well plate. The C_t_-values from the resulting plate were used to estimate the 2^-ΔCT^ with reference to *GAPDH* as internal reference. Further, the standard deviation and the standard error were calculated. The p-values were obtained based on Student’s *t*-test across 3 replicates. Each experiment has at least 3 independent biological replicates.

## Acknowledgements

We thank Prof. K. Veluthambi for *Agrobacterium* strains, PB-1 seeds, and binary plasmids and Dr. Dhandapani and many farmers for rice seeds. We thank central imaging facility (CIFF), genomics, electron microscopy, IT, radiation, greenhouse, and laboratory-kitchen facilities at the NCBS. We thank all the laboratory members for discussions and comments. This work was supported by NCBS-TIFR core funding, Department of Atomic Energy, Government of India, project identification no. RTI 4006 (1303/3/2019/R&D-II/DAE/4749 dated 16.7.2020) and grants (BT/IN/Swiss/47/JGK/2018-19; BT/PR25767/GET/119/151/2017) from Department of Biotechnology (DBT), Government of India. SR thanks Council of Scientific and Industrial Research (CSIR) for the funding. These funding agencies did not participate in the designing of experiments, analysis, or interpretation of data or in writing the manuscript.

## Author contributions

P.V.S. designed all experiments, discussed results, and wrote the manuscript with S.R. S.R generated the transgenic lines, performed most of the experiments and bioinformatic analysis. S.C. performed degradome and sRNA analyses. C.Y.H. performed molecular analysis and helped with protein analysis. All authors have read and approved the manuscript.

---

**Supplementary Figure 1.**
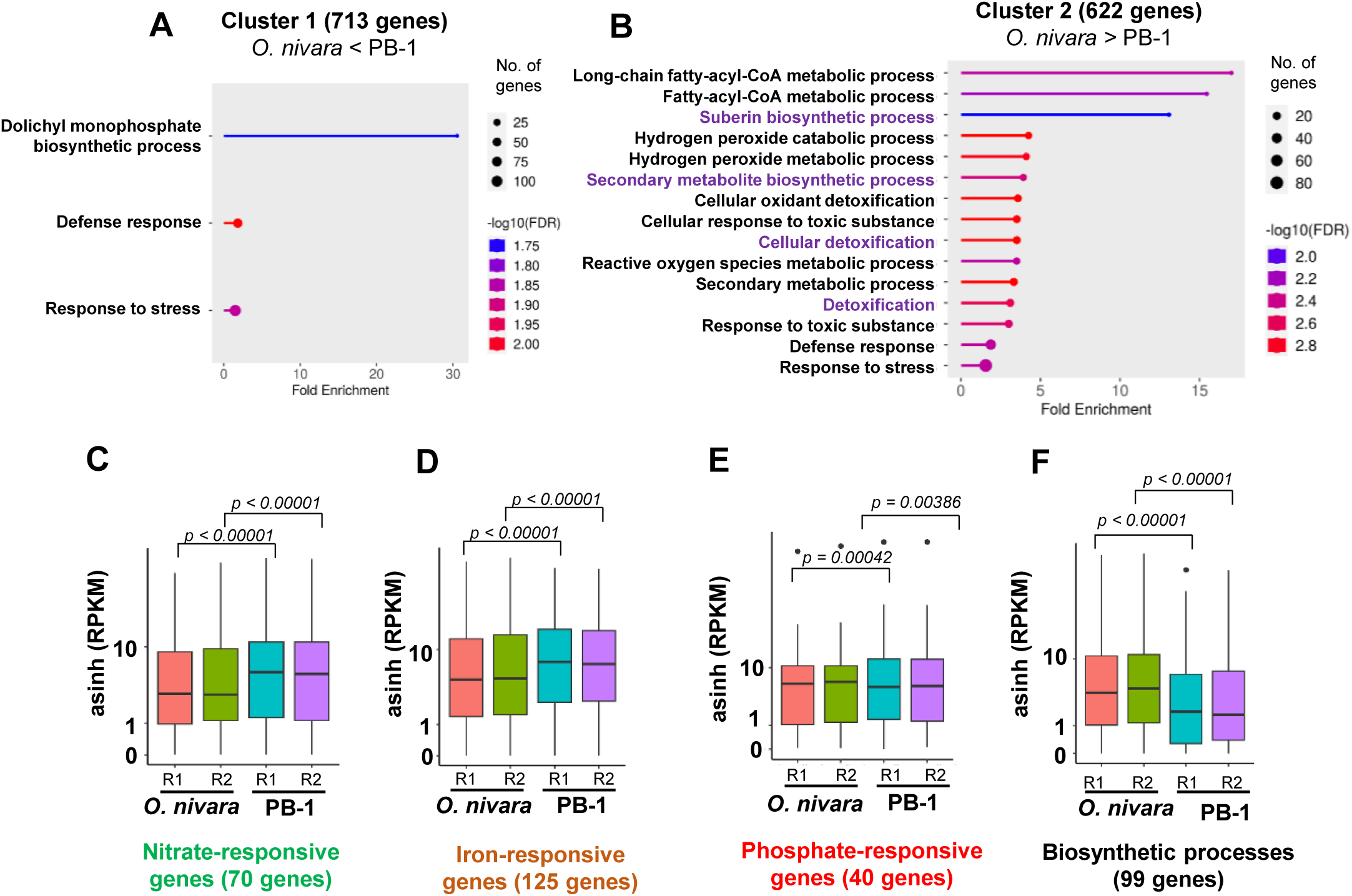
Wild and domesticated lines show differential expression of nutrient and stress responsive genes. (A) GO enrichment categories of Cluster 1 highly expressed DEGs. (B) Cluster 2 DEGs. PB-1 – 622 genes. (C) Box plot showing nitrate-responsive genes (n=70) and their expression across *O. nivara* and PB-1. (D) Box plot showing iron-responsive genes (n=125) and their expression across *O. nivara* and PB-1. (E) Box plot showing phosphate-responsive genes (40) and their expression. (F) Box plot showing biosynthetic process-associated genes (99) and their expression across *O. nivara* and PB-1. For C-F, asinh converted RPKM values were used for analysis. The boxes show median values and interquartile range. Whiskers show minimum and maximum values. Comparisons were made with two-sided Wilcoxon test (*p* < 0.05 was considered significant).

**Supplementary Figure 2.**
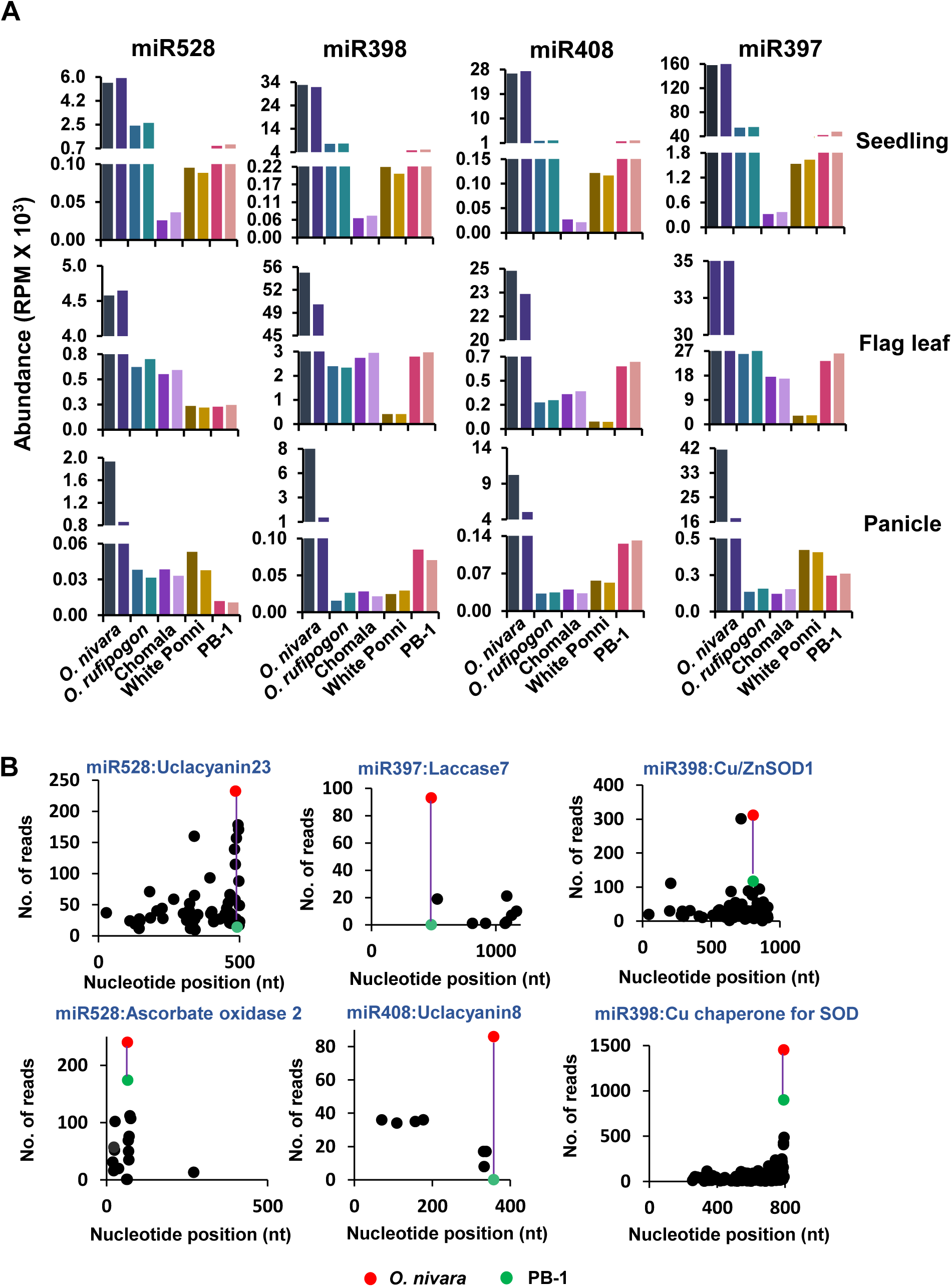
miRNA abundance and their targeting ability across rice lines. (A) miRNA abundance in 21-day old seedlings, flag leaf and panicle tissues. Abundance in reads per million (RPM) is plotted for each replicate. (B) Degradome T-plots showing targeting in wild and cultivated lines. Red dots-reads from *O. nivara,* green dots-reads from PB-1. Purple line indicates trend line in the degradome reads.

**Supplementary Figure 3.**
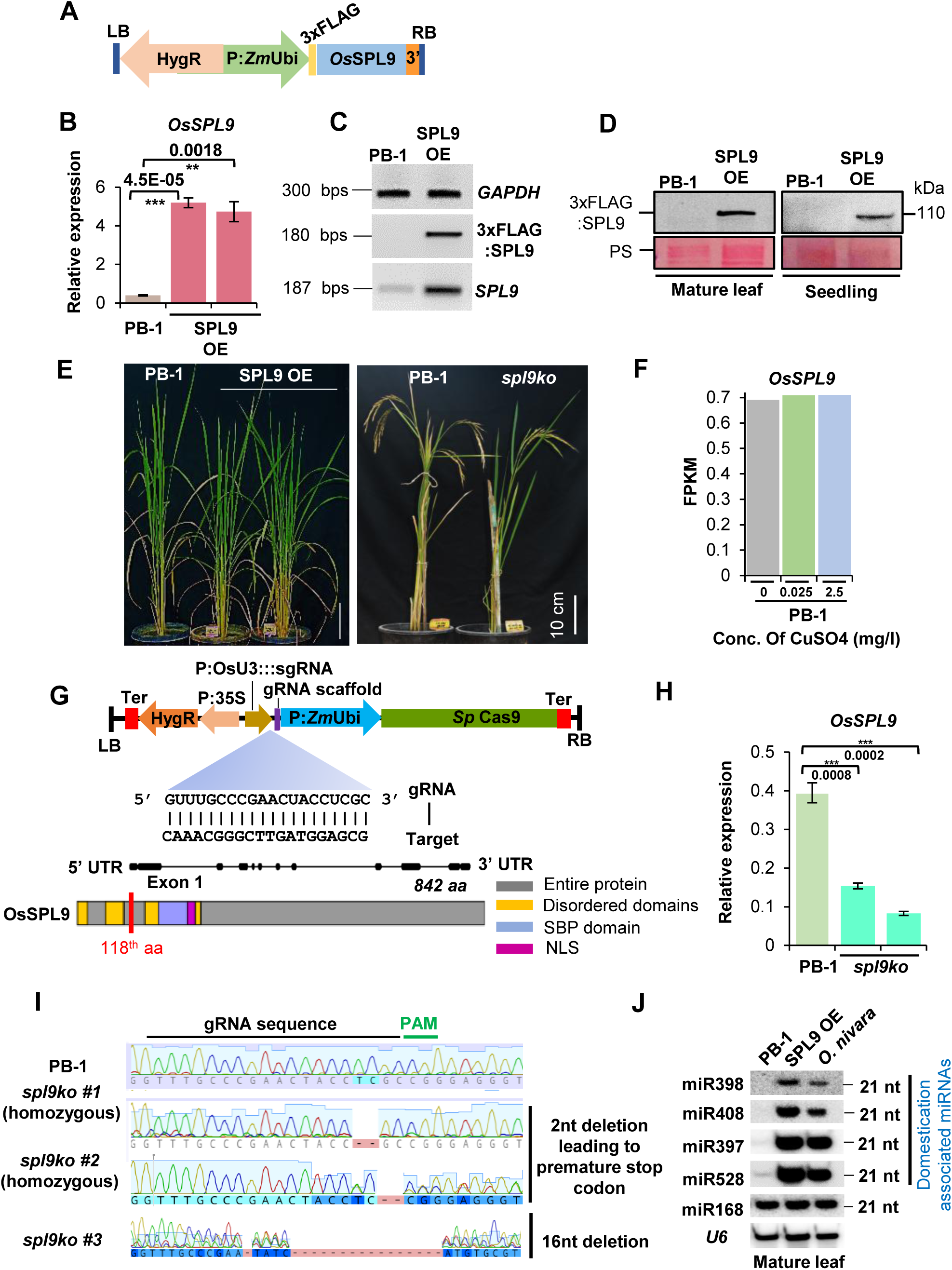
Generation and validation of SPL9 perturbed lines. (A) T-DNA map for SPL9 OE generation. (B) Relative expression of *OsSPL9* in lines expressing 3xFLAG:SPL9. (C) RT-PCR showing the levels of SPL9. *GAPDH* serves as internal control in B and C. (D) Western analysis showing the levels of 3xFLAG:SPL9 in mature leaf and seedling tissues. Ponceau (PS) serves as loading control. (E) Phenotype of the SPL9 mis-expression lines. Scale bar-10 cm. (F) Cuffnorm expression values (FPKM) of *OsSPL9* under Cu stress (mg/l) in PB-1. (G) T-DNA map showing gRNA targeting *OsSPL9* at exon 1 and the schematic representation of the transcript and domain structure of *OsSPL9*. The *spl9ko* leads to truncated protein. (H) Relative expression of *OsSPL9* in the *spl9ko* lines. *GAPDH* serves as control. (I) Chromatogram showing the region of editing in PB-1 and *spl9ko*. The gRNA and the adjacent PAM sequence are as mentioned. (J) Northern blotting showing the accumulation of domestication-associated miRNAs (miR528, miR397, miR408 and miR398) in mature leaf tissues. *U6* and miR168 serve as loading controls).

**Supplementary Figure 4.**
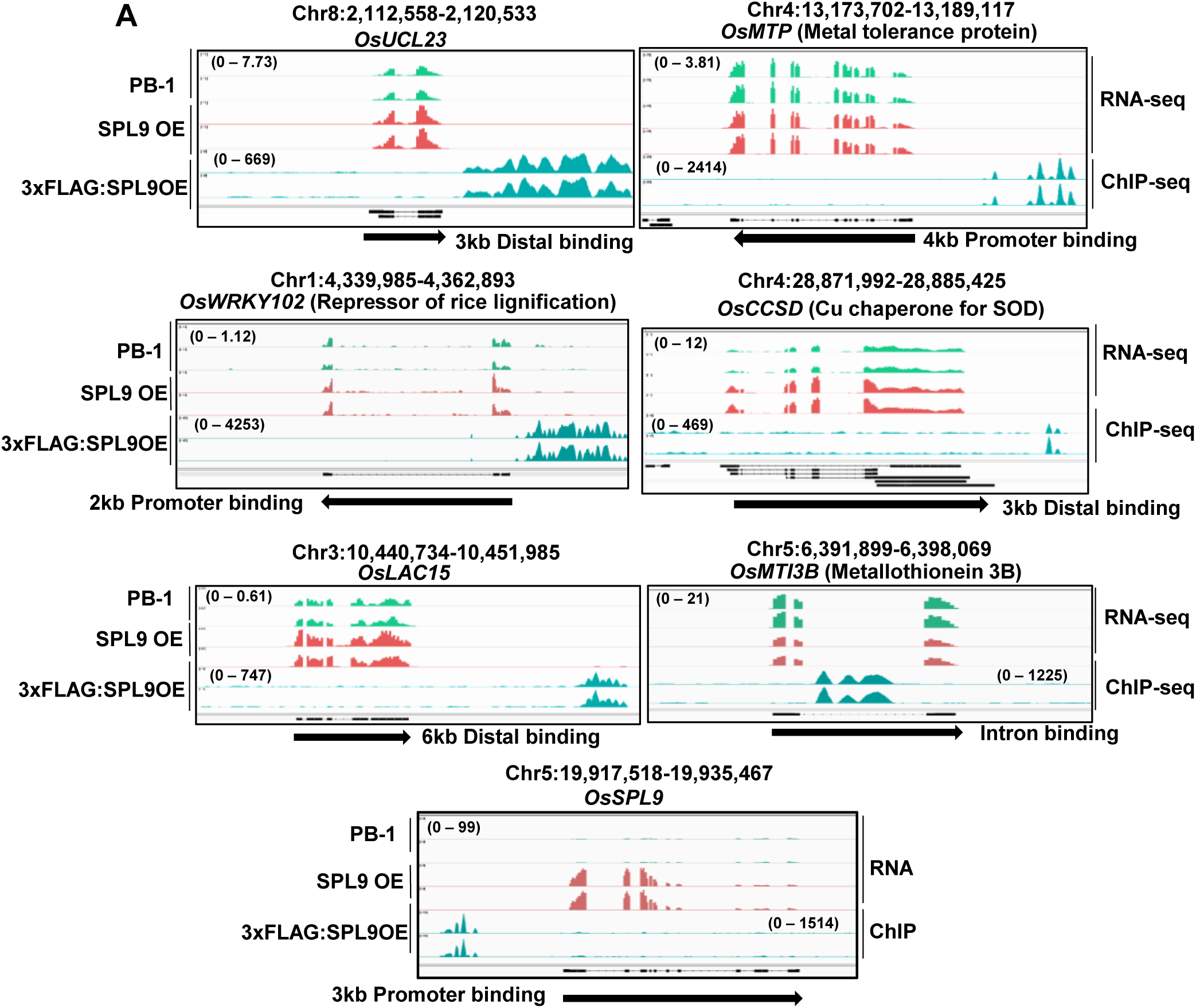
SPL9-TF binds to the promoters and genic regions of Cu-associated genes. (A) SPL9 peaks in upstream and downstream regions of several Cu-associated genes. Arrows indicate orientation of the gene and the panels indicate RNA-seq (top) and ChIP-seq (bottom). The track data ranges are mentioned in parentheses.

**Supplementary Figure 5.**
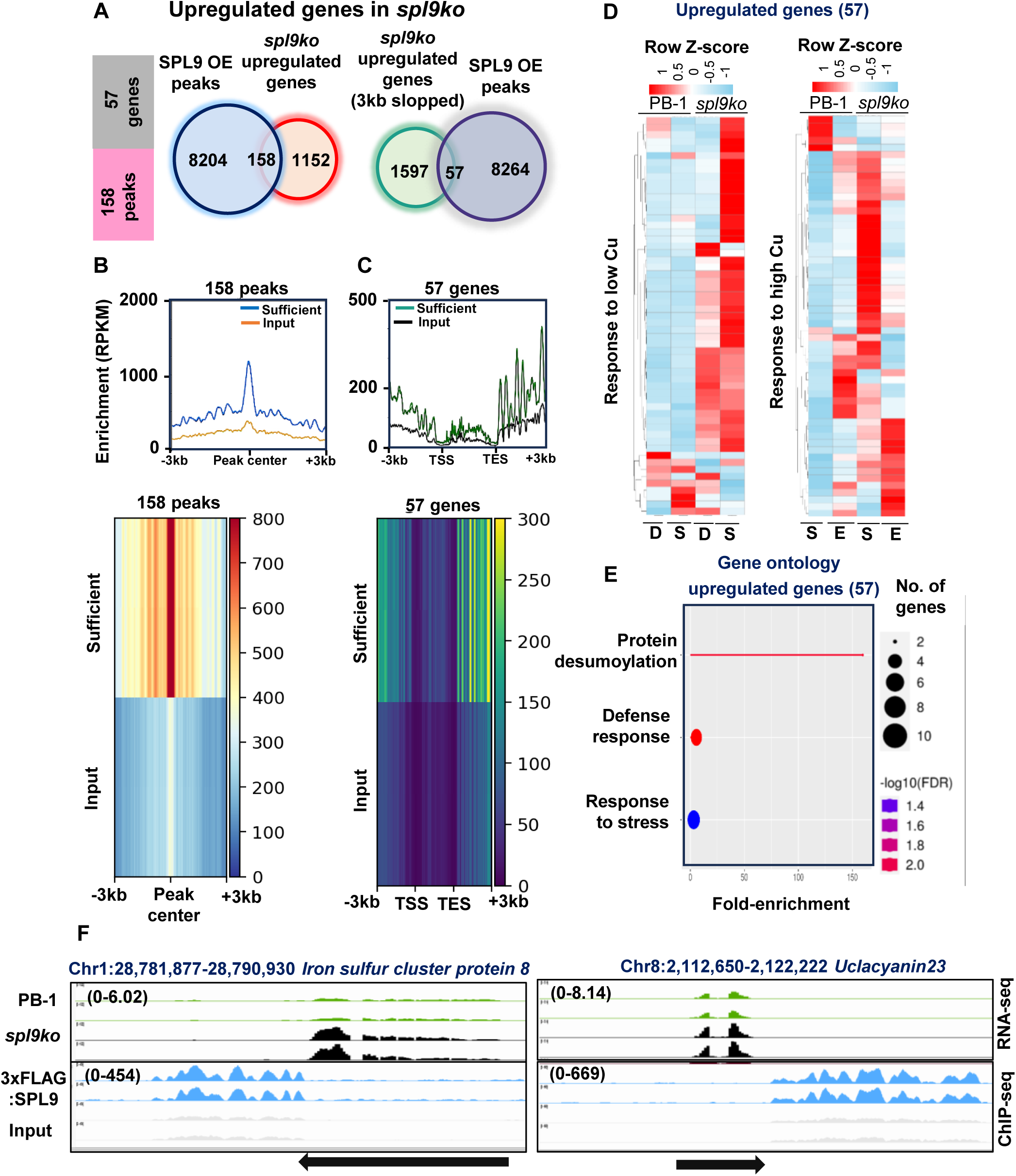
SPL9-bound upregulated genes shows response to Cu and have distinct peaks in their genomic regions. (A) Venn diagram for overlaps and cross-overlaps of SPL9 peaks in *spl9ko* upregulated genes. (B) Metaplot showing the overlaps and (C) cross-overlaps of SPL9 peaks on the upregulated genes. (D) Heatmap showing the SPL9-bound upregulated genes and their response to Cu (D – Cu-deficient, S – Cu-sufficient, E – Cu-excess). (E) Gene ontology categories of the SPL9-bound upregulated DEGs. (F) Genome browser screenshots of the SPL9-bound, upregulated DEGs in RNA-seq and ChIP-seq. Arrows indicate orientation of the gene and the panels indicate RNA-seq (top) and ChIP-seq (bottom). The track data ranges are mentioned in parentheses.

**Supplementary Figure 6.**
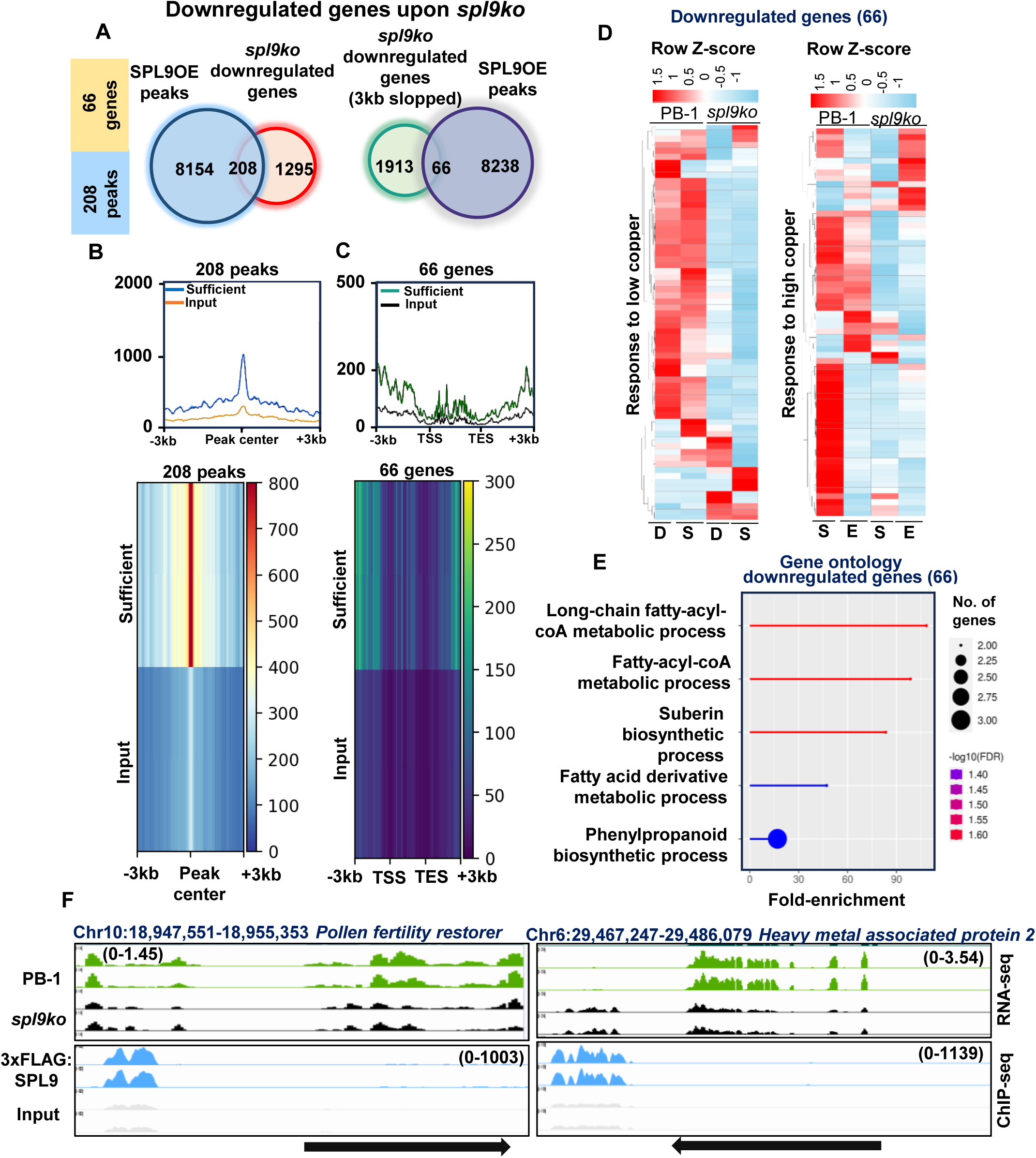
SPL9-bound downregulated genes show response to Cu and have distinct peaks. (A) Venn diagram for overlaps and cross-overlaps of SPL9 peaks on the *spl9ko* downregulated genes. (B) Metaplot showing the overlaps and (C) cross-overlaps of SPL9 peaks on the downregulated genes. (D) Heatmap showing SPL9-bound downregulated genes and their response to Cu. (E) Gene ontology categories of SPL9-bound downregulated DEGs. (F) Genome browser screenshots of SPL9-bound downregulated DEGs in RNA-seq and ChIP-seq. Other details are similar to Supplementary Figure 5.

**Supplementary Figure 7.**
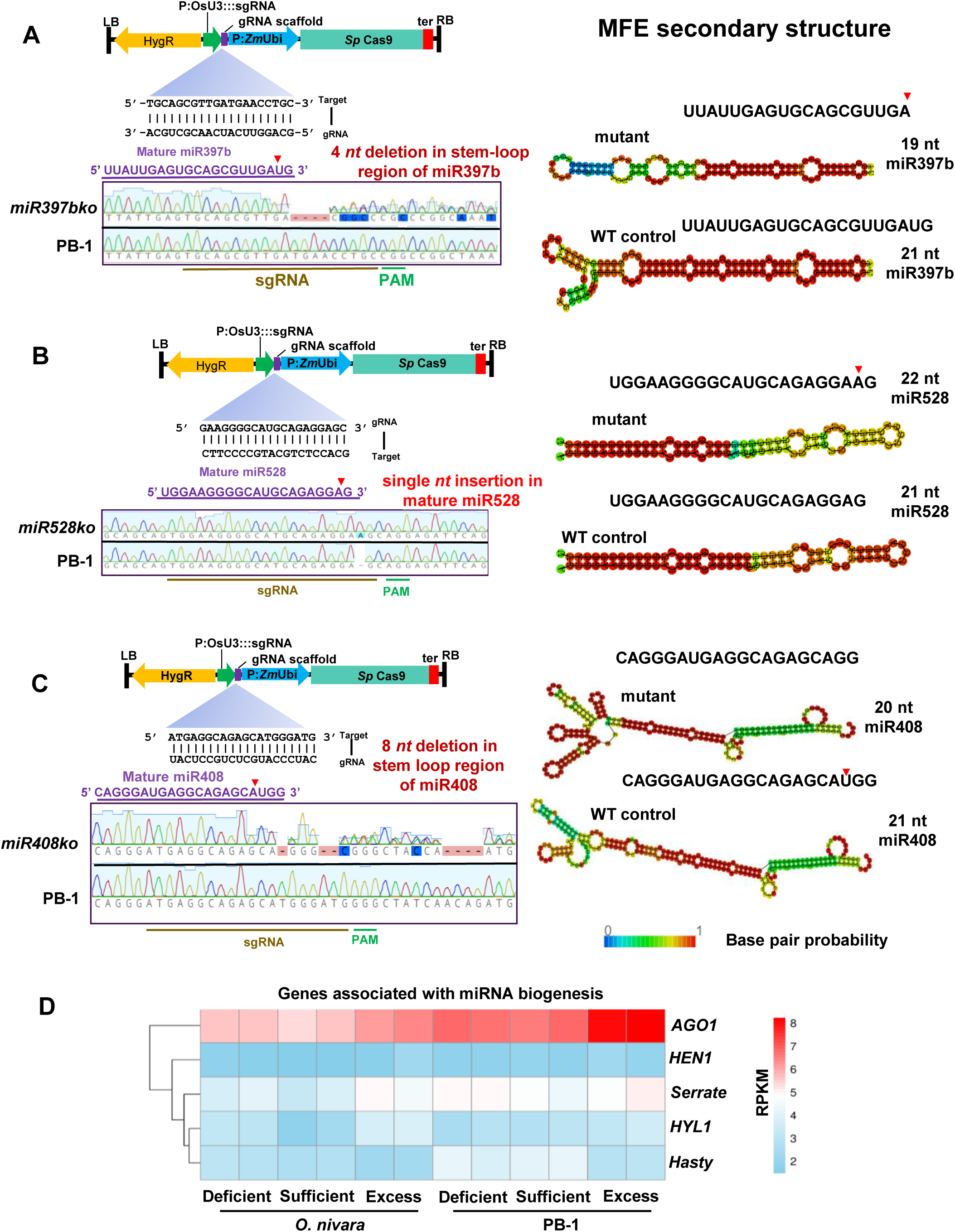
Schematic representation of miRNA ko mutants generated through CRISPR Cas9-mediated editing. T-DNA maps showing the gRNA targeting mature miRNA precursors of (A) miR397b, (B) miR528, and (C) miR408 are shown in the left. Minimum Free Energy (MFE) secondary structure of the mature miRNA of the mutants is shown on the right w.r.t to the WT control. (D) Heatmap showing the expression levels of miRNA biogenesis-associated genes across Cu stress conditions in *O. nivara* and PB-1.

**Supplementary Figure 8.**
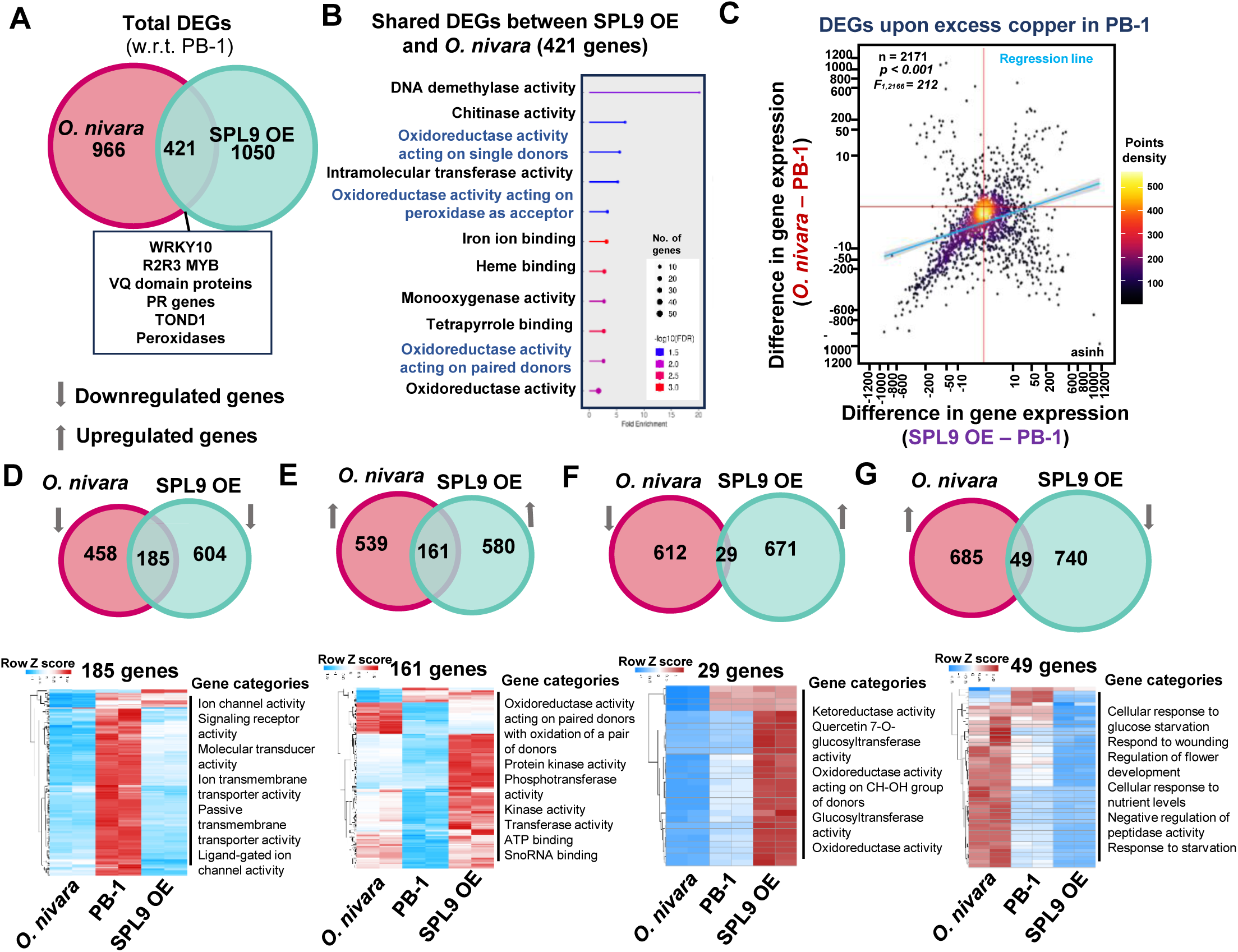
**Overlap of SPL9 OE transcriptome with *O. nivara.*** (A) Venn diagram showing overlap of total DEGs between *O. nivara* and PB-1. (B) Gene ontology showing the functions of the shared DEGs between *O. nivara* and PB-1. (C) Density scatter plot showing expression of DEGs upon excess Cu between *O. nivara* and SPL9 OE. Blue line-linear regression fit line. Grey shade represents 95 % confidence interval. Difference in gene expression is plotted with x-and y-axes are scaled to inverse sine hyperbolic function. F-test was used for statistical testing of the dependence of *O. nivara* and SPL9 OE. Points density are mentioned with colour gradient. (D) Venn diagram showing the overlap of downregulated genes in SPL9 OE and *O. nivara* and the heatmap for the expression of 185 shared genes. (E) Overlap of upregulated genes in SPL9 OE and *O. nivara* and the heatmap showing the expression of 161 shared genes. (F) upregulated genes in SPL9 OE and downregulated genes in *O. nivara* and the heatmap showing the expression of 29 shared genes. (G) Downregulated genes in SPL9 OE and upregulated genes in *O. nivara* and the heatmap for 49 shared genes. The GO category of each gene set is mentioned on the right side.

**Supplementary Figure 9.**
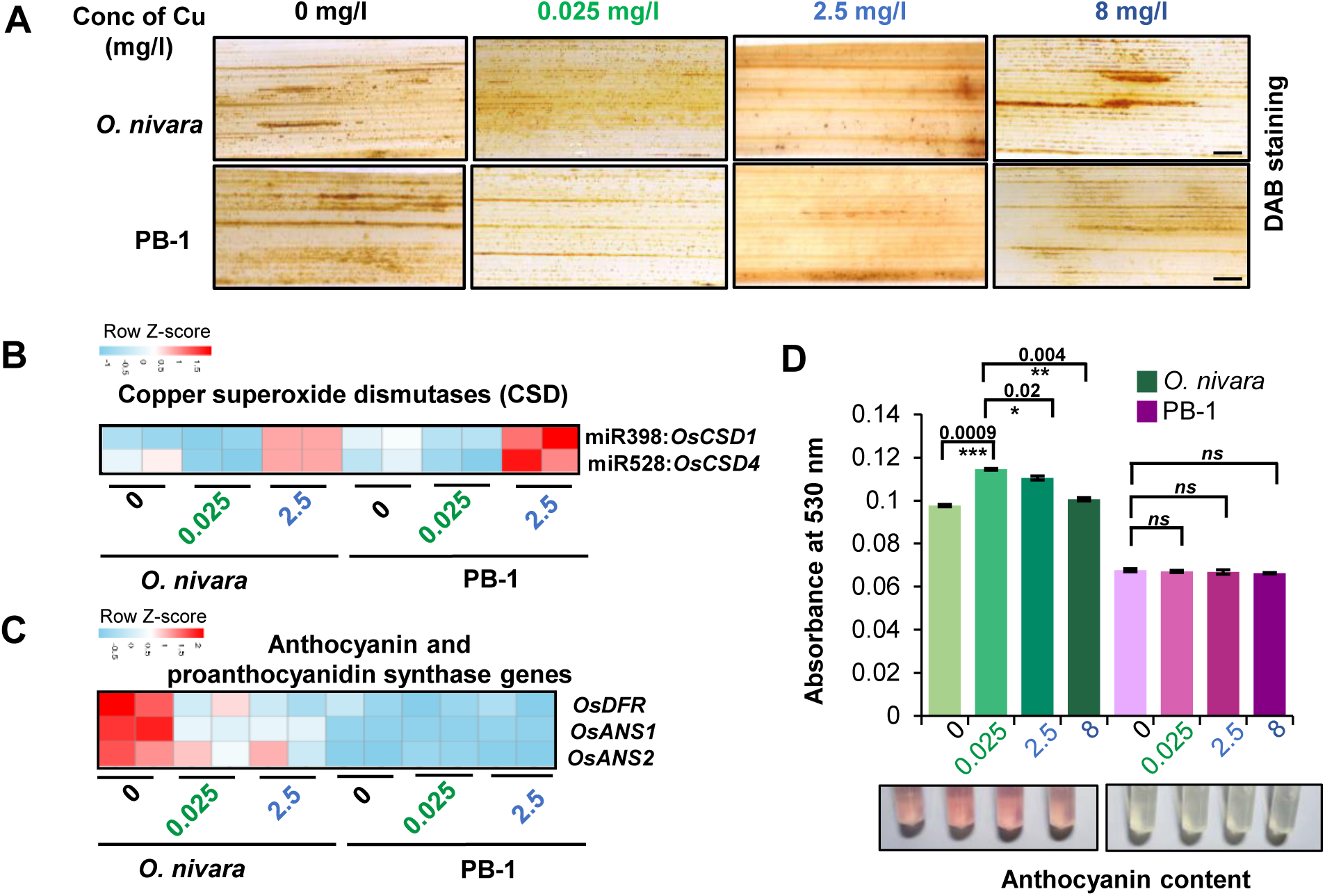
The SOD activity and anthocyanin accumulation changes in wild and cultivated rice lines across varying Cu concentrations. (A) Leaves of *O. nivara* and PB-1 upon DAB staining. (B) Heatmap showing expression of copper superoxide dismutase (CSD) genes in *O. nivara* and PB-1 under Cu stress. (C) Heatmap showing expression of anthocyanin and proanthocyanidin synthase genes in *O. nivara* and PB-1. (D) Anthocyanin accumulation in *O. nivara* and PB-1 under Cu stress. Two-tailed Student’s *t*-test was used for statistical comparison. (*) *p*-value < 0.05, (*ns*) non-significant.

**Supplementary Table 1:**
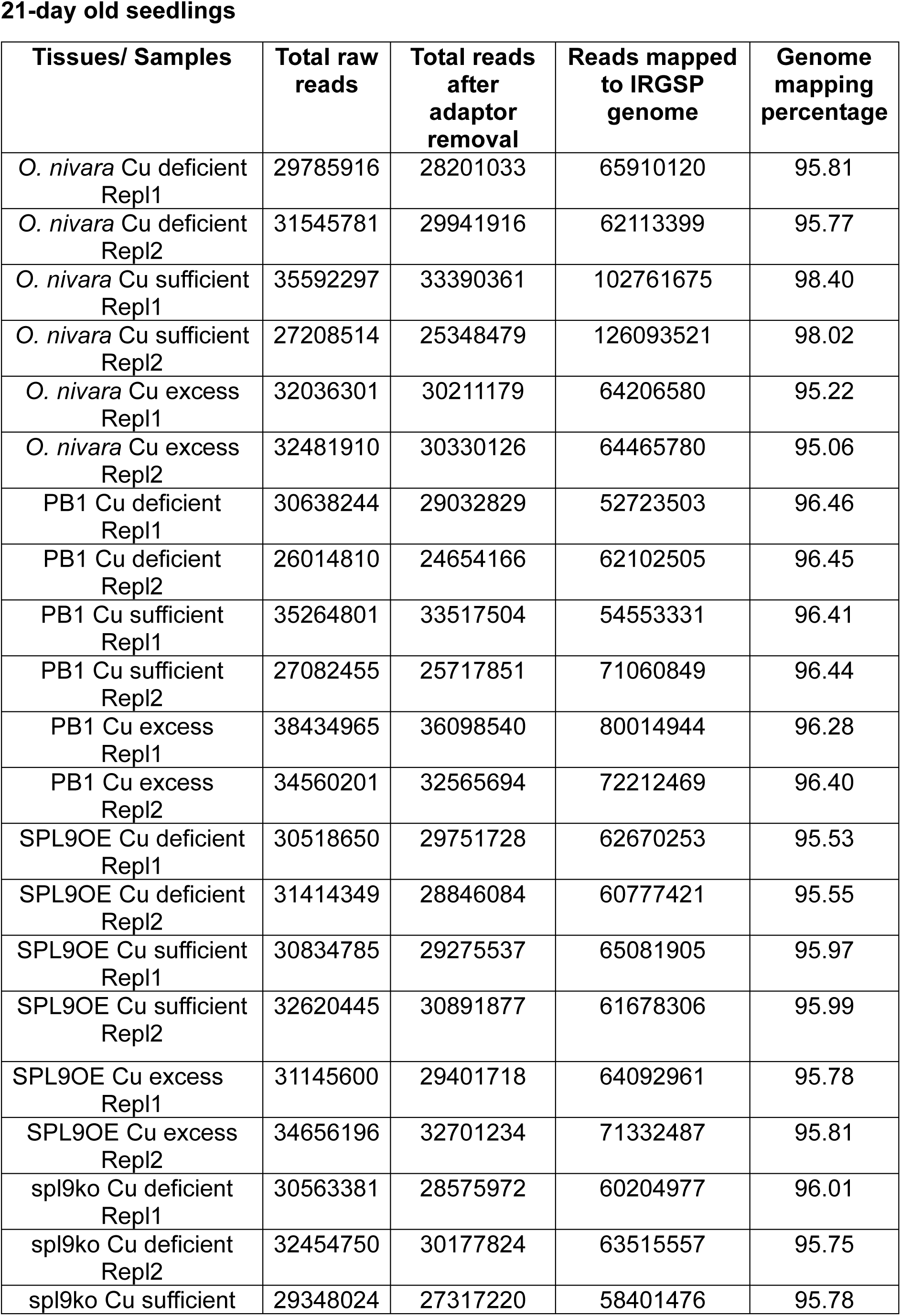
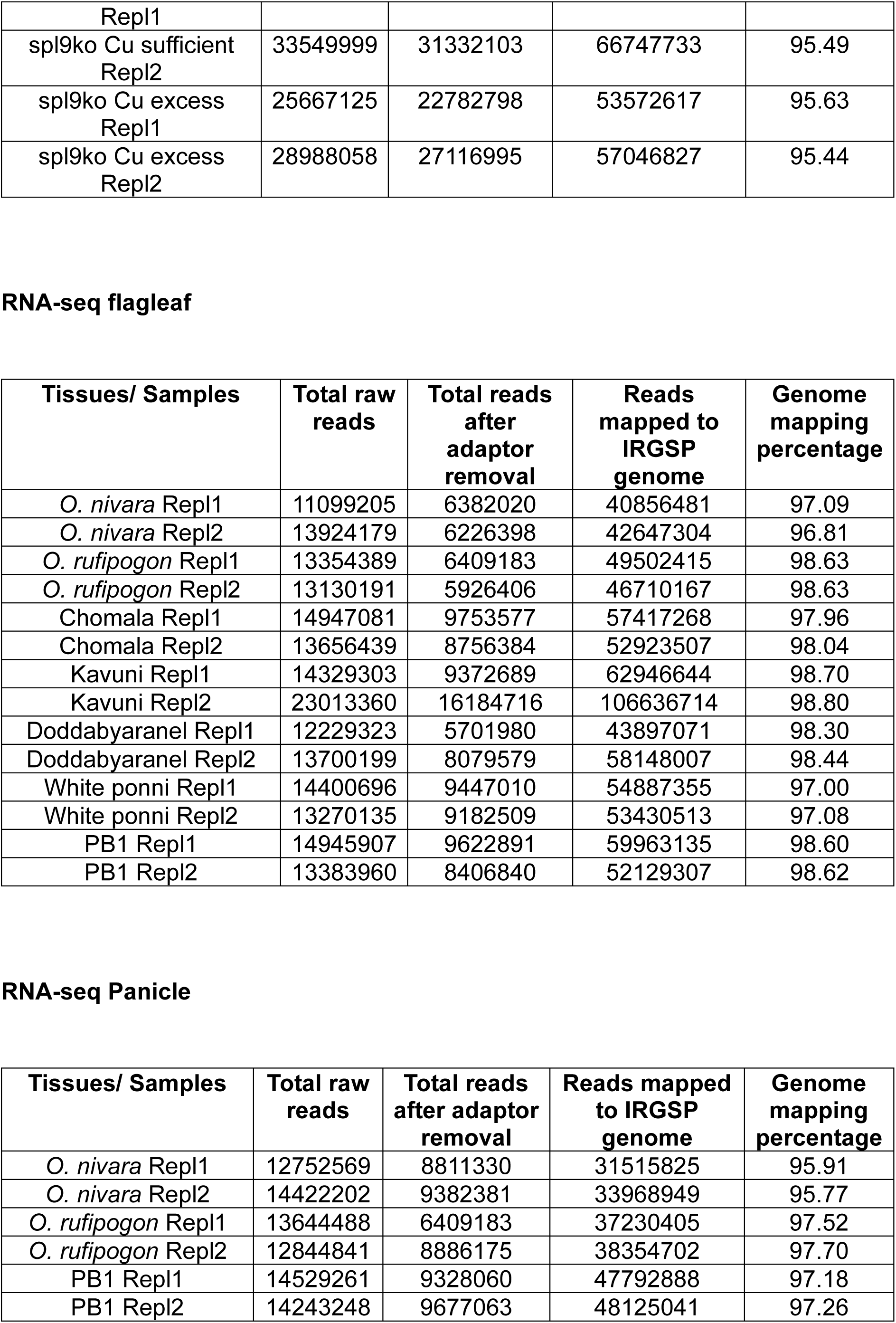
Details of library statistics used in RNA-seq datasets.

**Supplementary Table 2:**
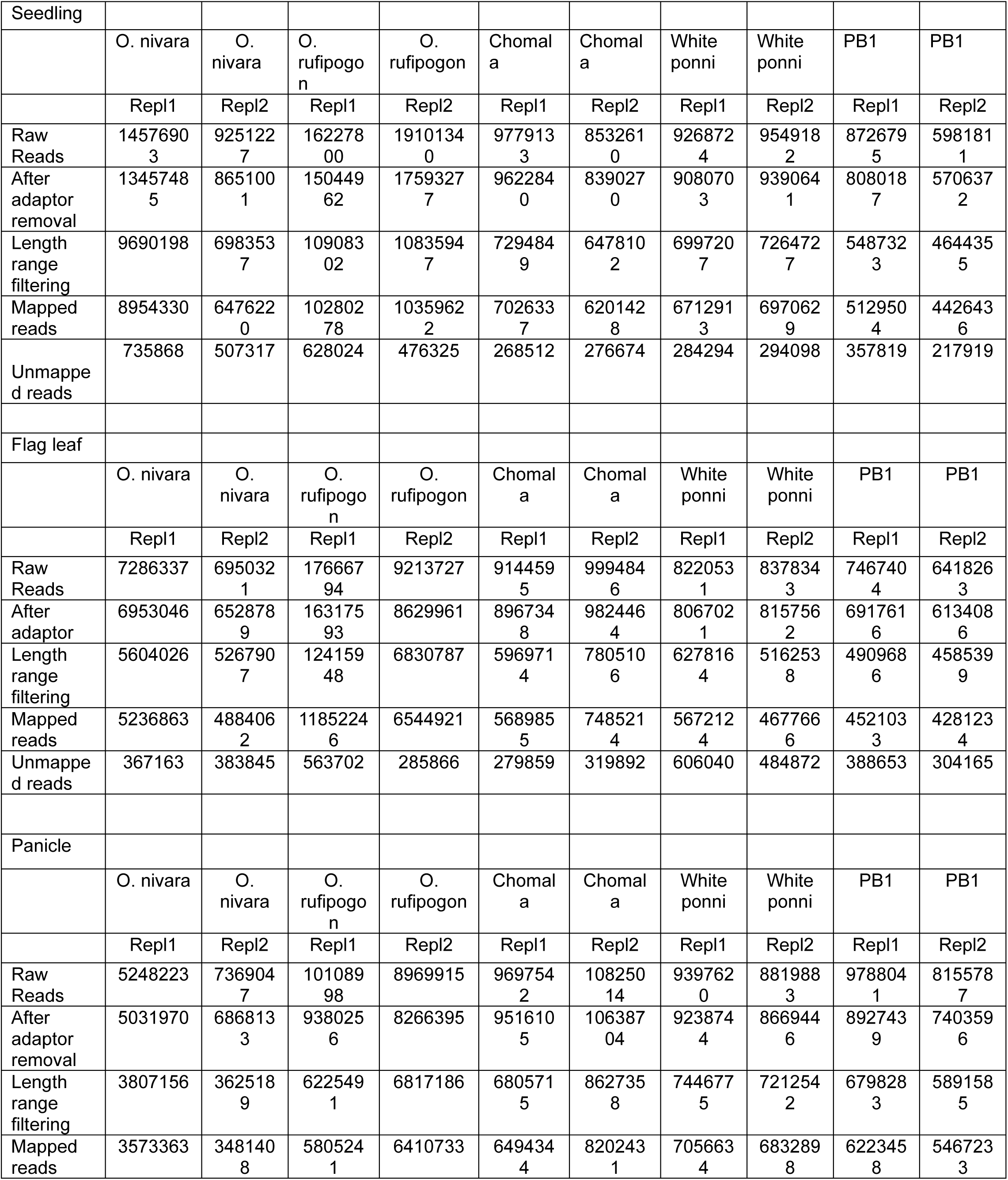
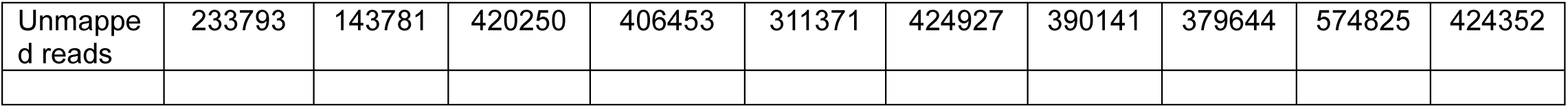
Details of library statistics used in sRNA-seq datasets.

**Supplementary Table 3:** Details of library statistics used in ChIP-seq datasets.

21-day old seedlings
| Tissues/ Samples | Total raw reads | Total reads after adaptor removal | Reads mapped to IRGSP genome | Genome mapping percentage |
| --- | --- | --- | --- | --- |
| SPL9OE Cu deficient Repl1 | 20849645 | 20815435 | 56877484 | 67.56 |
| SPL9OE Cu deficient Repl2 | 21314392 | 21308021 | 63111722 | 70.65 |
| SPL9OE Cu sufficient Repl1 | 21656971 | 21501063 | 57752890 | 66.78 |
| SPL9OE Cu sufficient Repl2 | 20150860 | 20135878 | 58458446 | 69.73 |
| SPL9OE Cu excess Repl2 | 22927151 | 22880072 | 64040030 | 68.28 |
| SPL9OE Cu excess Repl1 | 21744430 | 21739138 | 64325608 | 70.91 |
| Input_Repl1 | 30832564 | 30772335 | 29476820 | 95.79% |
| Input_Repl2 | 29989331 | 29939706 | 28634335 | 95.64% |

**Supplementary Table 4:** Transformation summary.

| **Binary plasmid** | **Calli used for transformation** | **Calli showing positive response** | **No. of regenerated calli** | **Regeneration percentage** | **Shoot per regenerated calli (n=5)** |
| --- | --- | --- | --- | --- | --- |
| pVHS20<br>SPL9OE | 250 | 15 | 10 | 4 | Three |
| pRGEB32<br>spl9ko | 280 | 8 | 5 | 1.78 | Two |
| pRGEB32<br>528ko | 300 | 7 | 2 | 0.6 | Two |
| pRGEB32<br>408ko | 320 | 4 | 2 | 0.625 | Three |
| pRGEB32<br>398ako | 260 | 12 | 3 | 1.15 | One-Two |
| pRGEB32<br>397bko | 350 | 10 | 5 | 1.42 | Two-Three |

**Supplementary Table 5:**
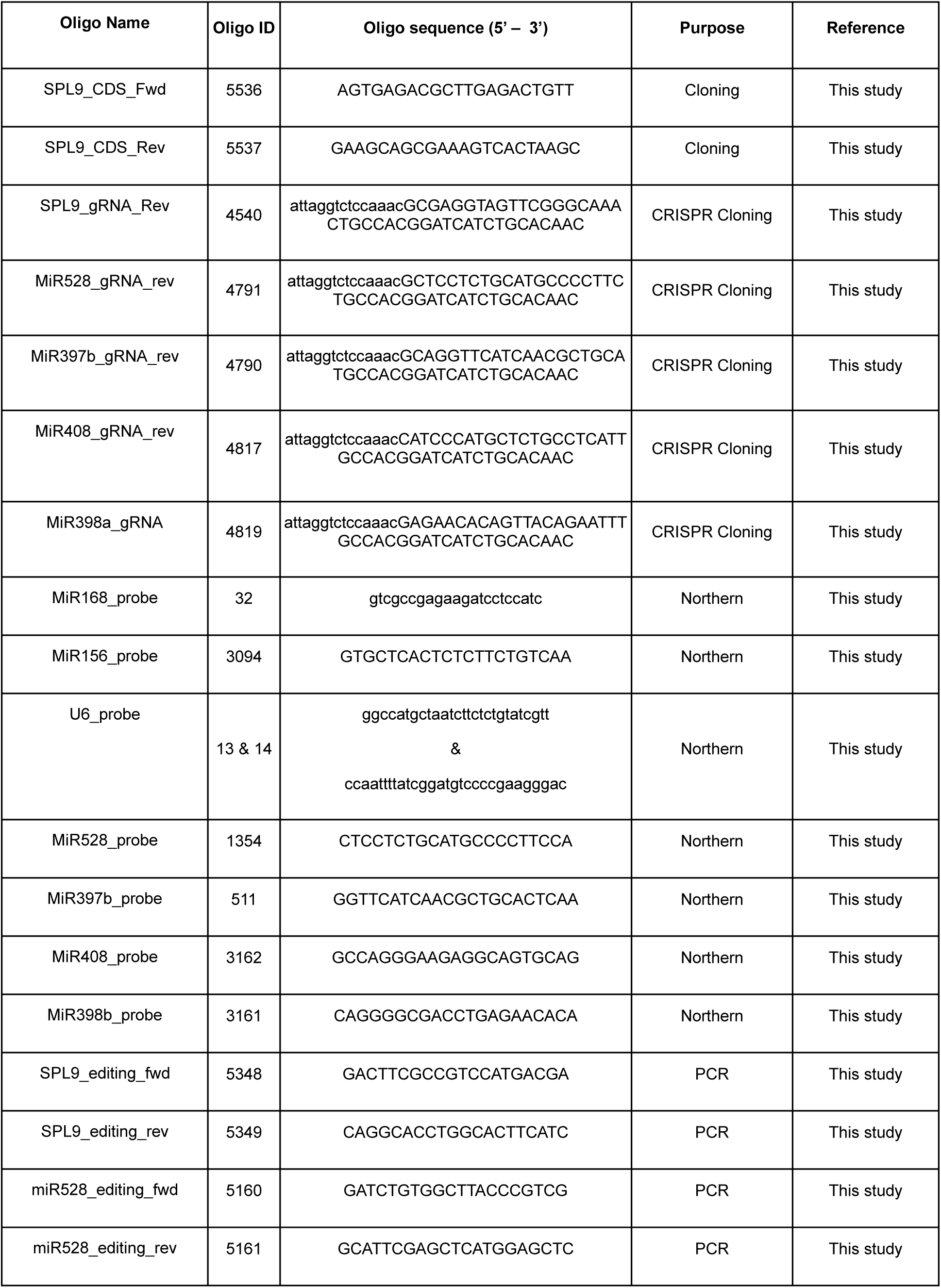
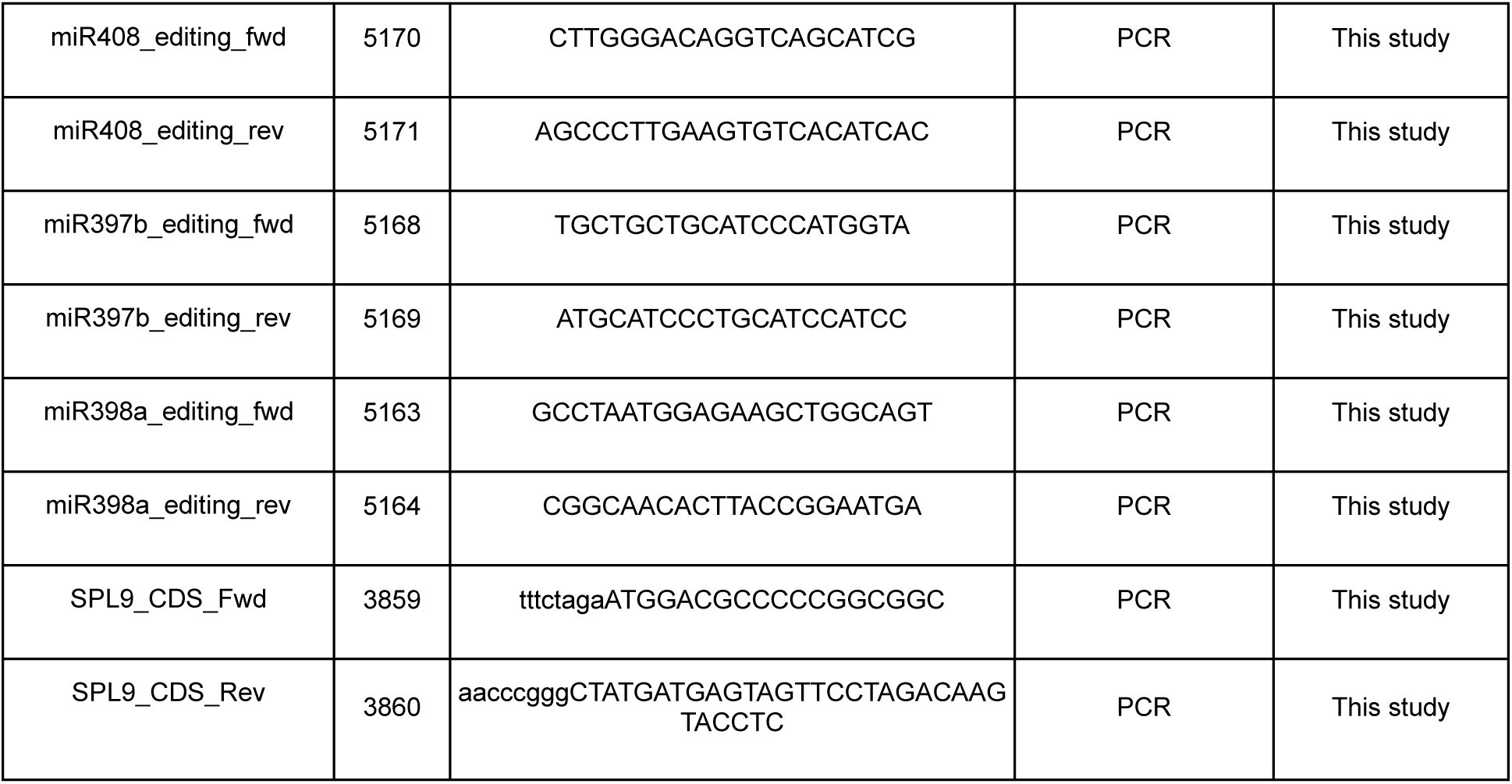
List of oligos and probes used in this study.

**Supplementary Table 6:** Details of the published dataset used in this study (Wang *et al.,* 2012).

|  | Expression (RPM) |  |  |  |  |
| --- | --- | --- | --- | --- | --- |
|  | *O. rufipogon* |  |  | *O. sativa japonica* |  |
|  | leaf | root | developing seed | leaf | developing seed |
| **miR408** | **75.5** | 37.3 | 3.9 | **14.4** | 25.2 |
| **miR528** | **10815.5** | 90.4 | 428.5 | **310.3** | 1226.7 |
| **miR397a** | **5.7** | 45.3 | 1.9 | **19.3** | 6.7 |
| **miR397b** | **0.1** | 0.0 | 0.0 | **0.0** | 0.0 |
| **miR398b** | **2.5** | 0.0 | 0.3 | **12.9** | 0.7 |

